# Actuation of CRP activating region 3 by acetylation modulates *V. cholerae* sugar utilization and virulence

**DOI:** 10.64898/2026.01.27.701997

**Authors:** Renato E.R.S. Santos, Jacob A. Gibson, Pallabi Basu, Michael J. Gebhardt, Chris Akut, William P. Robins, John J. Mekalanos, Simon L. Dove, Paula I. Watnick

**Affiliations:** Division of Infectious Diseases, Boston Children’s Hospital, 300 Longwood Avenue, Boston, MA 02115, USA; Department of Pediatrics, Harvard Medical School, 25 Shattuck Street, Boston, MA 02115, USA; Biological and Biomedical Sciences Program, Harvard Medical School, 25 Shattuck Street, Boston, MA 02115, USA; Department of Microbiology and Immunobiology, University of Iowa, USA; Department of Microbiology, Harvard Medical School, 25 Shattuck Street, Boston, MA 02115, USA

**Keywords:** *V. cholerae*, cAMP receptor protein, lysine acetylation, virulence, acetate switch, small RNA

## Abstract

The cyclic AMP receptor protein or CRP is a global regulator of bacterial metabolism that activates transcription of genes required for utilization of alternative carbon sources in response to the second messenger cAMP, which is synthesized in the setting of glucose scarcity. CRP activates transcription through contact with RNA polymerase at three sites termed activating regions (ARs) 1-3. AR3 was previously reported to be functional only when CRP K52 was mutated to a neutral residue and to be essential for transcription only in the absence of AR1 and AR2. Multiple proteomic studies have reported acetylation of CRP K52. This post-translational modification is predicted to activate AR3. To probe the role of K52 acetylation (K52QAc) and AR3 at the genome level, we used ChIP-seq and RNA-seq analysis to compare WT CRP with a CRP K52Q mutant that mimics CRP K52Ac. We report that CRP K52Q binds to hundreds of new sites on the chromosome, resulting in increased abundance of known as well as previously unknown transcripts. These transcripts increase uptake and metabolism of dietary sugars such as maltose and galactose, repress acetate consumption, and augment virulence gene expression. We attribute the repression of acetate consumption to a novel small RNA, CrbZ, which is positively regulated by CRP K52Q in LB broth and by WT CRP specifically in minimal medium containing maltose. This study highlights the role of post-translational modifications in molding the CRP regulon to optimize pathogen metabolism and virulence gene expression in the human intestine in response to nutritional cues.

**Significance statement:** As a model in the field of bacterial transcription, the structure and function of the cAMP receptor protein (CRP), a global transcription regulator, has been exhaustively investigated. These studies have established three activating regions (ARs) where CRP contacts RNA polymerase, of which only two were thought to participate in transcription activation by native CRP. Here we provide evidence that post-translational acetylation of *V. cholerae* CRP lysine 52 actuates AR3, enabling occupancy of hundreds of novel CRP binding sites and the transcription of genes encoding novel small RNAs. These changes alter virulence gene expression, promote utilization of dietary carbon sources, and delay acetate uptake. We propose that acetylation of CRP K52 engages AR3, thus optimizing *V. cholerae* fitness in the human intestine.

## Introduction

The cyclic AMP receptor protein (CRP) is a conserved global regulator of bacterial transcription (1). To regulate transcription, CRP dimers complexed with the second messenger cyclic adenosine monophosphate (cAMP) bind to the chromosome at two inverted repeats with the consensus sequence ATGTGA separated by a 6 bp spacer (Fig 1A) (2).

**Figure 1:**
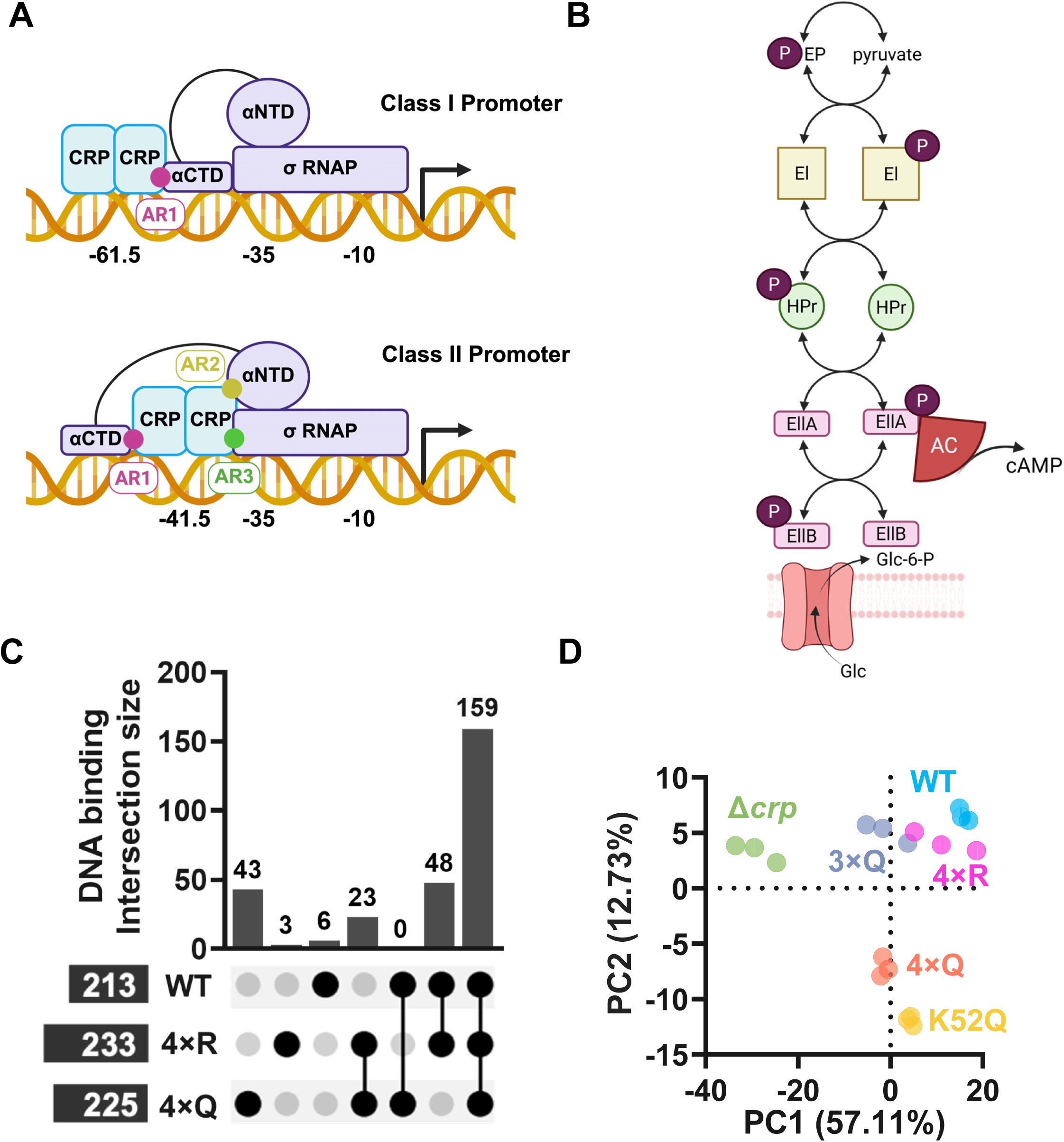
Point mutants designed to mimic unmodified and acetylated CRP modulate DNA occupancy, gene transcription, and virulence. (A) Illustration showing the interaction of CRP activating regions (AR) 1-3 with RNA polymerase (RNAP) subunits α and σ at Class l and Class ll promoters. *Created in BioRender. Watnick, P.* (*2026*) https://BioRender.com/2vpegk4. (B) Schematic showing the phosphoenolpyruvate phosphotransfer cascade. Enzyme l (El) is phosphorylated by phosphoenolpyruvate (PEP). This phosphate is passed to Histidine protein (HPr) and from there to enzymes llA and B (EllA, EllB). Enzyme llC (EllC) is an integral membrane transporter that mediates tandem sugar transport and phosphorylation by EllB. *Created in BioRender. Watnick, P.* (*2025*) https://BioRender.com/Sm3al62. (C) UpSet plot showing common and unique CRP DNA occupancy sites identified by ChIP-seq for WT, 4×Q, and 4×R CRP. Occupancy sites were defined by an enrichment factor (EF) > 4 for at least 2 of 3 biological replicates. Shared occupancy events are indicated by connecting lines. (D) Principal Component Analysis of differentially expressed genes (-1 ≤log_2_(FC)≥ 1 and FDR < 0.05) based on RNA-seq analysis of the indicated strains cultured in LB medium and harvested at mid-exponential growth phase. Biological triplicates are shown.

CRP activates uptake and catabolism of many sugars when glucose is unavailable, and glucose transport is intimately tied to generation of cAMP. The phosphoenolpyruvate phosphotransferase system (PTS), a regulatory multi-component phosphotransfer cascade that terminates in phosphorylation of the incoming sugar, mediates transport of a variety of sugars (Fig 1B) (3). PTS transport of some sugars depends on CRP, while transport of others such as glucose does not. The PTS cascade begins with phosphorylation of Enzyme I (EI) by phosphoenolpyruvate. This phosphate is passed to Histidine protein (HPr) and from there to sugar-selective enzymes llA and B (EllA and B). Sugar transport through the Enzyme llC results in passing of phosphate from Enzyme llB directly to the incoming sugar. Enzymes ll A, B, and C may be independent proteins or domains within a single multidomain protein. In *V. cholerae*, the sugar specificities of many Ell components have been determined (4, 5).

When sugars transported by the PTS are abundant, sugar-specific PTS proteins exist principally in their unphosphorylated state, while they are phosphorylated when these sugars are absent. Both the phosphorylated and unphosphorylated states of these proteins regulate cell physiology by direct protein-protein interactions. PTS transport plays a key role in regulation of transcription by CRP because phosphorylated glucose-specific Enzyme llA interacts directly with adenylate cyclase to increase intracellular levels of cAMP (6, 7). Thus, when glucose is scarce, CRP is active. Association of cAMP with the N-terminus of CRP increases RNA polymerase binding and transcription, which activates hundreds of genes including many that transport and metabolize alternative carbon sources (8). In many pathogens, CRP also regulates virulence gene expression (9–13).

In initial studies of *Escherichia coli* CRP, two classes of promoters were identified that make distinct contacts with the α-subunit of RNA polymerase (Fig 1A). At class I promoters, the consensus binding sequence of CRP is centered at -61.5 bp with respect to the transcription start site (TSS). This results in contact between the downstream CRP monomer and the C-terminal domain of the α subunit (α-CTD) of RNA polymerase (RNAP). CRP residues 156-164, which interact with the α-CTD are termed activating region 1 (AR1) and are essential for transcription activation at class l promoters (14).

At class ll promoters, the CRP consensus binding sequence is centered at -41.5 bp with respect to the TSS. Here the α-CTD of RNAP wraps around the CRP dimer via a flexible loop, and the AR1 contact forms between the upstream CRP monomer and the α-CTD of RNAP. At class ll promoters, two additional activating regions located in the downstream CRP monomer (AR2 and AR3) have been identified. AR2, which consists of CRP residues 19, 21, 96, and 101, contacts the N-terminal domain of the α subunit of RNAP and is essential for transcription at class ll promoters (15, 16). In contrast, AR1 is only essential at a subset of class ll promoters. CRP AR3 is defined by amino acid residues 52-58 which contact the σ-subunit of RNAP (17, 18). AR3 was reported to activate transcription at class ll promoters only when K52 is changed to a neutral amino acid and transcription activation by AR1 or AR2 is diminished by mutation (19, 20). This body of work led to the conclusion that CRP AR3 is dispensable for transcription activation by native CRP at class ll promoters (14).

The genome of the diarrheal pathogen *Vibrio cholerae* encodes a CRP homolog. Of the 210 amino acids that comprise *V. cholerae* CRP, 201 are identical to those of *Escherichia coli* CRP including the residues defining AR3. CRP is crucial for *V. cholerae*’s response to the aquatic environment, mammalian hosts, environmental hosts, and *Vibrio*-specific phages through its regulation of metabolism, quorum sensing, horizontal gene transfer, biofilm formation, and virulence (21–32). Interestingly, although CRP represses many virulence determinants including the toxin co-regulated pilus, cholera toxin, the RTX toxin, and biofilm formation under laboratory conditions, a Δ*crp* mutant, in which these virulence factors are highly expressed, is massively defective for colonization of both the mammalian and zebra fish intestines (27, 33). Two explanations are that *V. cholerae* CRP is essential for optimal utilization of nutrients to promote growth in the human intestine and/or that CRP does not actually repress virulence gene expression *in vivo*.

We previously reported that *V. cholerae* CRP is membrane-associated under specific growth conditions and sequesters the dual function transcription factor peptidase A to the membrane (34). In these studies, we found that K52 was succinylated. A proteomic study detailing the acetylome of the closely related species *Vibrio parahemolyticus* as well as proteomic studies of purified *E. coli* CRP have reported acetylation of K52 (35, 36). Because acetylation neutralizes the positive charge of K52, this post-translational modification is predicted to engage AR3.

To better understand the impact of K52 acetylation on the function of *V. cholerae* CRP, we defined the regulon of a CRP mutant with K52 substituted for Q to mimic acetylation. Here we report that a CRP K52Q substitution generates hundreds of novel chromosomal binding sites, some of which result in increased abundance of putative small RNAs (sRNAs). This significantly reshapes the regulon of CRP leading to increased transcription of genes that encode virulence factors and sugar metabolism genes and repression of those involved in acetate utilization. We show that the latter are regulated by the novel sRNA CrbZ, which is in turn positively regulated by CRP K52Q in cells grown in LB broth and by wild-type (WT) CRP in cell grown in minimal medium containing maltose. Although CRP K52Q reverses inhibition of the *V. cholerae* virulence factors by WT CRP, this mutant has no competition advantage in the infant mouse intestine. We hypothesize that CRP K52 is acetylated *in vivo* resulting in engagement of AR3 to optimize *V. cholerae* fitness in the mammalian intestine.

## Results

### An amino acid substitution that mimics CRP K52 acetylation modulates DNA occupancy, gene expression, and virulence without altering CRP membrane association

We previously reported that, when *V. cholerae* was cultured in defined medium supplemented with sucrose, CRP was found principally in the membrane fraction (34). When lysines 22, 26, 35, and 52 were mutated to Q (4XQ) to mimic acetylation, CRP relocated to the cytoplasm, leading to the hypothesis that post-translational modifications dictate the subcellular localization of CRP. In *V. cholerae* LB broth cultures, approximately 26% of CRP was found in the membrane (Fig S1). To determine if similar residues might determine the subcellular localization of CRP in LB broth, we tested chromosomal 4XQ and 4XR mutants designed to mimic acetylated and deacetylated forms of CRP, respectively. A V5 affinity tag was added to the C-terminus of WT CRP and all mutants to facilitate subsequent analysis. The 4XQ mutant was found principally in the cytoplasmic fraction, while the 4XR mutant was more abundant in the membrane fraction (Fig S1).

Because a sizeable amount of CRP was cytoplasmic for all *V. cholerae* strains cultured in LB broth, we hypothesized that ChIP-seq under these conditions might elucidate the impact of these substitutions independent of differences in membrane association. Importantly, the V5 tag used for immunoprecipitation had only a small effect on gene expression (Fig S2). ChIP-seq analysis was carried out on biological triplicates of WT, 4XR, and 4XQ mutants cultured in LB broth to mid-log phase. Enrichment in DNA occupancy was measured relative to an untagged control and a minimum of 4-fold enrichment (EF≥4) in each of three biological replicates was considered significant (Table S1). As shown in Fig 1C, all three alleles were significantly enriched at the majority of chromosomal binding sites, proving that all alleles are functional. WT CRP and the 4XR mutant were most similar with both showing enrichment at an additional 48 sites. In contrast, while WT CRP and the 4XR mutant bound very few sites uniquely, the 4XQ substitution enhanced binding at multiple novel chromosomal locations. Twenty-three sites were common between the 4XR and 4XQ CRP mutant alleles, indicating a unique role for lysine in inhibiting binding at these sites.

To investigate the functional significance of the additional 4XQ binding sites, we undertook RNA-seq analysis of exponential phase cultures of WT CRP as well as 4XR, 4XQ, and Δ*crp* mutants (Table S2). Because the 4XQ mutant includes K52Q, which is predicted to engage AR3, we included K52Q and 3XQ mutants (K22Q, K26Q, and K35Q) in our analysis (15, 16, 18, 37). WT CRP and all mutants were expressed at comparable levels, grew at comparable rates, and did not alter acetylation at other sites (Fig S3). Furthermore, the K52Q substitution did not alter CRP membrane association (Fig S1). We defined significantly differentially expressed genes as those having at least a 2-fold change in expression and a false discovery rate of 0.05 or less. Table 1 shows the numbers of total and gene-associated transcripts that increased and decreased in abundance relative to a Δ*crp* mutant. WT CRP and the 4XR mutant had similar numbers of differentially regulated genes, while the 3XQ mutant differentially regulated far fewer genes than WT CRP, and the K52Q mutant differentially regulated more genes than any of the other alleles. The 4XQ mutant regulated fewer genes than WT and K52Q and more than 3XQ, indicating a phenotype intermediate between the K52Q and 3XQ mutants.

**Table 1:** Genes differentially regulated relative to the. Δ*crp* mutant in strains expressing the indicated V5-tagged CRP isoforms from the chromosomal locus. **Table 1: Transcripts differentially regulated relative to the** Δ*crp* mutant in strains expressing the indicated chromosomal V5-tagged CRP isoforms.

| vs $\Delta crp$ | All transcripts | | Coding transcripts only | |
| --- | --- | --- | --- | --- |
|  | Up | Down | Up | Down |
| WT | 1405 | 1035 | 639 | 508 |
| 4XR | 1455 | 981 | 658 | 485 |
| 4XQ | 1137 | 754 | 525 | 341 |
| 3XQ | 800 | 660 | 395 | 309 |
| K52Q | 1606 | 1320 | 729 | 566 |

To further explore the RNA-seq data, we carried out principal component analysis. The WT, 4XR and 3XQ CRP transcriptomes clustered together, while the 4XQ and K52Q transcriptomes were separate and closer to each other (Fig. 1D). Taken together, the ChIP-seq and RNA-seq data suggest that the K52Q substitution, and by extension post-translational acetylation of K52, has a large impact on transcription regulation by CRP. We, therefore, focused on the role of the CRP K52Q substitution in chromosomal occupancy and gene regulation.

We first performed a ChIP-seq experiment comparing chromosomal binding of WT CRP and the K52Q protein (Table S3). As shown in Fig 2A, using an enrichment factor of 4 (EF>4) as a threshold, WT CRP associated with a mean of 66 chromosomal sites, while CRP K52Q bound to 263 sites, an increase of almost 4-fold. Considering sites that reached the threshold of EF>4 in 2 of 3 replicates, only 2/3 of WT binding sites were shared by K52Q (Fig 2B). We then categorized the positions of the chromosomal binding sites of WT CRP and the K52Q mutant with respect to putative TSSs using a previous comprehensive report of *V. cholerae* TSSs as a reference (38). Putative promoters were defined as 200 bp upstream of a TSS (Fig 2C). The 5 untranslated region (5 -UTR) was considered to extend from the TSS to the start of the coding sequence (CDS) of a gene. The majority of WT CRP binding sites were within promoters (Fig 2C). While very few WT CRP binding sites were located outside promoters, almost half of the K52Q mutant binding sites were either in the 5 -UTR or CDS of genes.

**Figure 2.**
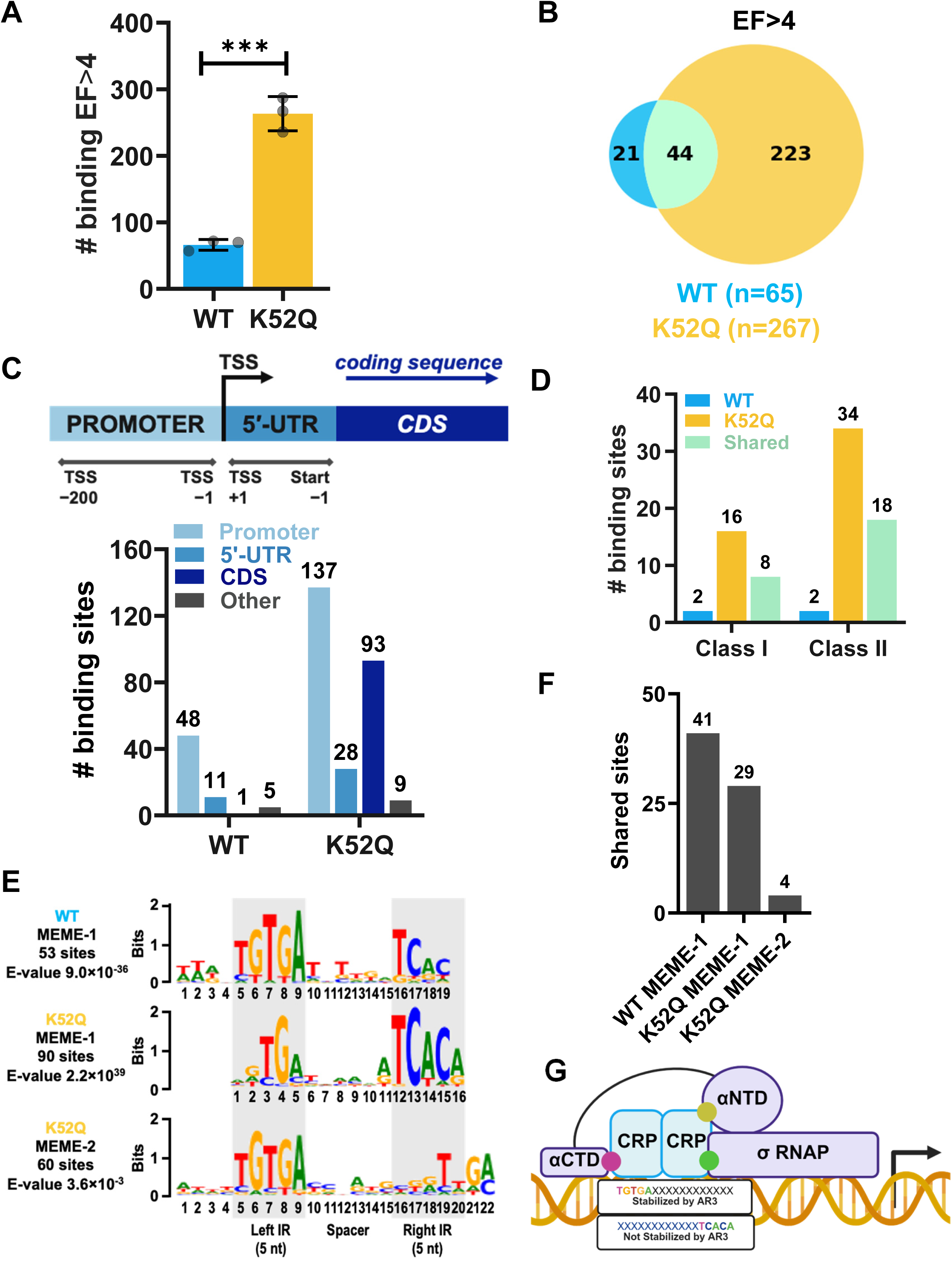
Many chromosomal binding sites unique to K52Q CRP are distinguished by their operator sequence and position within coding sequences. (A) Total number of sites where WT CRP and CRP K52Q binding reach EF >4. Bars indicate the average of three biological replicates. Error bars represent the standard deviation. A student’s t test was used to calculate significance. (B) Venn diagram showing total numbers of binding sites that are unique and shared between WT CRP and a CRP K52Q mutant. These numbers reflect peaks that were present in 2 out of the 3 biological replicates. EF>4 was used as a threshold. (C) Map showing how positions of WT and K52Q binding sites were categorized, and enumeration of WT and K52Q CRP chromosomal binding sites that are in the regions indicated. (D) Numbers of class I and class ll promoters identified for WT CRP and CRP K52Q. Class l promoters were defined as within -52 to -70 of a TSS, while class ll promoters were defined as within -51 to -32 of TSS. (E) Consensus motifs discovered by MEME (Multiple Em for Motif Elicitation, https://meme-suite.org/meme/). The number of sites and the corresponding E-values are indicated. E-values are derived from the log-likelihood ratio (LLR) statistic used by MEME’s expectation-maximization (EM) algorithm and represent the expected number of motifs with an equal or higher LLR occurring by chance under the background model. (F) Graph showing numbers of chromosomal binding sites used to build MEME 1 and 2 that are shared by WT CRP and the CRP K52Q mutant or unique to the CRP K52Q mutant. (G) Schematic showing orientation of Meme ll operators in which association of the unbound CRP K52Q monomer is stabilized by an interaction of AR3 with σ^70^. The green circle represents AR3. The arrow indicates the direction of transcript elongation. Created in BioRender. Watnick, P. (2025) https://BioRender.com/2vpegk4.

In a CRP K52Q mutant, AR3 is predicted to form an additional contact with the σ subunit of RNAP at Class ll promoters. In the few promoters previously studied, K52 and AR3 did not participate in transcription activation by native CRP at Class ll promoters (18, 37). Nevertheless, we were curious whether Class ll promoters might be over-represented in K52Q binding sites. We defined Class l promoters by the location of a CRP binding site within -69 to -52 bp of a TSS and class ll promoters as a binding site within -50 bp to -32 bp of the TSS. Based on these definitions, we identified binding at more class ll than class I promoters for both WT CRP and CRP K52Q (Fig 2D). While the majority of WT CRP binding sites at Class l and Class ll promoters were shared by CRP K52Q, CRP K52Q had approximately twice as many Class l and Class ll binding sites as WT CRP, suggesting that CRP K52Q does not show a preference for binding at Class ll promoters.

### CRP K52Q tolerates more degeneracy in operator sequences than WT CRP

We questioned whether the operator sequences bound by WT CRP were distinct from those bound by CRP K52Q. Based on 53 binding sites, Meme analysis of WT CRP yielded an operator consensus sequence that was similar to that previously defined for *E. coli* and *V. cholerae* CRP (Table S4, Fig 2E) (2, 27, 39). Meme analysis of the CRP K52Q binding sites yielded two consensus sequences. One of these (Meme l), based on 90 binding sites, was closely related to that of WT CRP. The second consensus sequence (Meme ll), based on 60 binding sites, displayed degeneracy in one half of the inverted repeat, suggesting that, at some sites, binding of one half of the CRP K52Q dimer to the chromosome may be adequate to stabilize the interaction with DNA.

We reasoned that WT CRP and the K52Q mutant might share some of the binding sites on which their respective consensus sequences were based. In fact, out of the 53 sites used to identify WT Meme l, 41 were shared with CRP K52Q (Fig 2F). In contrast, only 29 of the 90 sites used to identify CRP K52Q Meme l and only 4 of the 60 sites used to identify CRP K52Q Meme ll were shared.

The σ^70^ subunit of RNAP, which interacts with CRP AR3, binds to the -35 and -10 elements within promoters where it stabilizes the open complex of DNA known as the transcription bubble, which is essential for transcript elongation (40). Promoter escape after synthesis of the first 11 nucleotides of the mRNA was reported to coincide with release of σ^70^ from RNAP, at least partly due to displacement by the growing mRNA transcript (41). However, recent *in vitro* studies suggest that the σ^70^ subunit of RNAP can be retained throughout transcript elongation and that the interaction of σ^70^ with sequences similar to σ^70^ binding sites, which are found throughout the genome, may induce pausing of the elongation complex (42–44). We, therefore, hypothesized that, if the Meme ll operator was oriented appropriately, CRP K52Q binding to the DNA, even at intragenic sites, could be stabilized by the interaction of CRP K52Q AR3 with σ^70^ (Fig 2G). Sixteen Meme ll sites were in promoters, while 44 were in coding sequences (Table S4). We examined whether each Meme ll consensus sequence was oriented appropriately to allow an interaction between AR3 of the unbound CRP K52Q and σ^70^. We found that 38/60 or approximately 2/3 of the Meme ll sites were appropriately oriented, while the other 1/3 were not. This suggests that binding of CRP K52Q at Meme ll sites is independent of an interaction with σ^70^. Last, we examined whether CRP K52Q is functional at Meme ll operator sites. CRP K52Q enrichment at Meme ll sites positioned within genes was not correlated with an effect on transcript abundance. In contrast, 18 Meme ll binding sites were within promoters. Of these, 9 were correlated with either positive or negative changes in transcript abundance as compared with WT CRP. Taken together, these observations suggest that the CRP K52Q substitution stabilizes binding to Meme ll operator sites independent of an interaction with σ^70^ and further that, when positioned within promoters, binding of CRP K52Q to Meme ll sites can alter transcription.

### CRP K52Q activates gene transcription at Class ll promoters

To determine whether CRP K52Q was more likely than WT CRP to increase transcript abundance, we plotted transcript abundance against the presence of WT CRP binding only, CRP K52Q and WT binding, or CRP K52Q binding only (Fig 3A). In each case, both increases and decreases in transcript abundance were observed, suggesting that CRP K52Q does not serve exclusively as an activator of transcription. We then questioned whether CRP K52Q was more likely to activate transcription at Class ll promoters than Class l promoters. At the few binding sites shared by WT CRP and CRP K52Q within promoters meeting the definitions of Class l or Class ll, gene expression was equally likely to be positively or negatively regulated at both types of promoters (Fig 3B). In contrast, at sites bound only by CRP K52Q, transcription activation was more likely than repression at Class ll but not Class l promoters (Fig 3C). This is consistent with engagement of AR3 by CRP K52Q and, possibly, CRP K52Ac, at Class ll promoters. Interestingly, in the majority of cases, CRP K52Q enrichment at both Class l and Class ll promoters had no effect on transcription. This suggests either that these promoters were incorrectly classified or that they are not active during growth in LB broth. Last, we explored the multitude of CRP K52Q binding sites that are within CDSs to determine whether these affect expression of the corresponding gene. As shown in Fig 3D, only 20 out of the 93 CRP K52Q binding sites in CDSs altered transcription. Twelve of these positively regulated transcription while 8 negatively regulated transcription.

**Figure 3.**
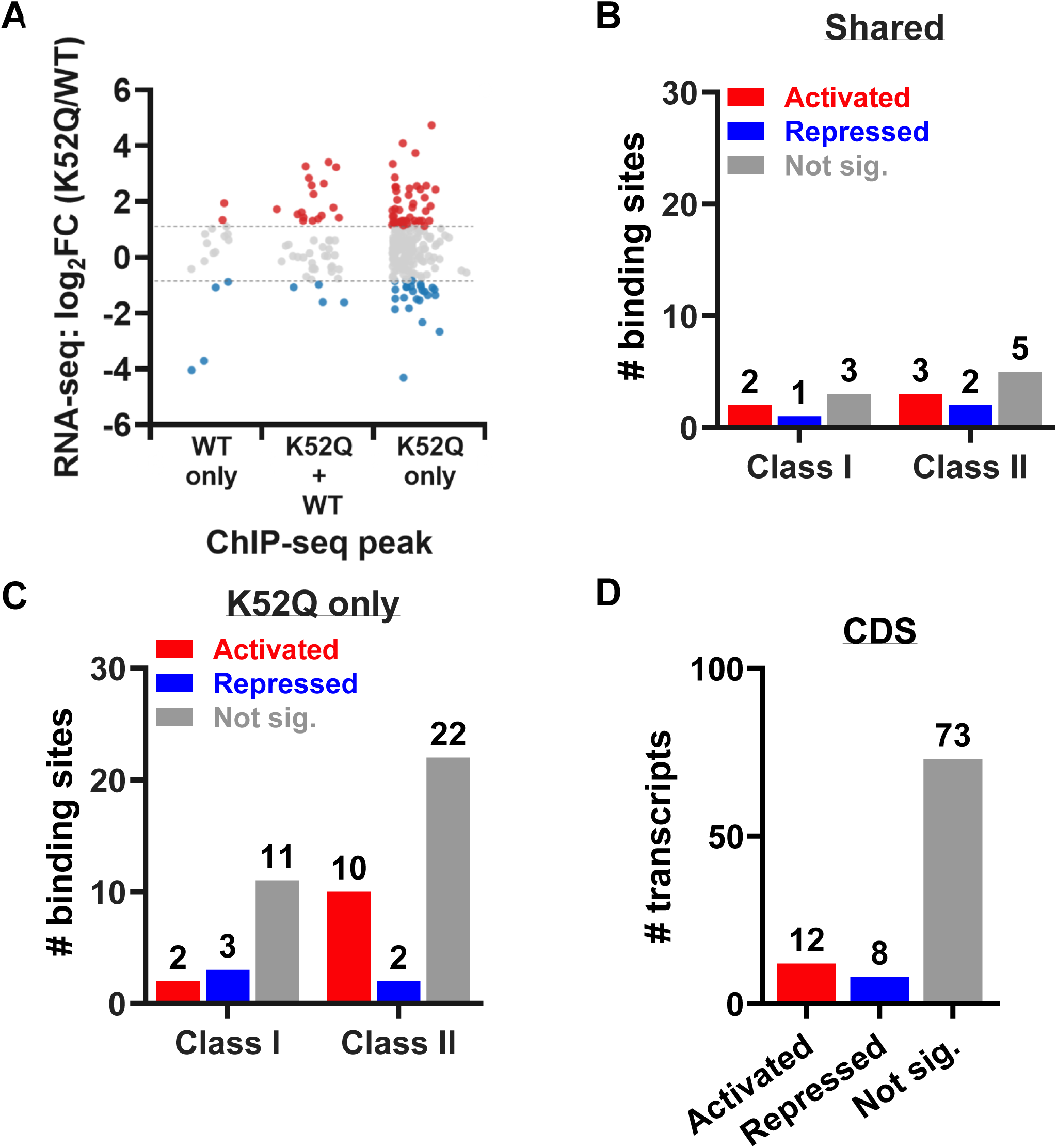
CRP K52Q regulates transcription at many new class ll promoters and intragenic binding sites. (A) Scatter plot of differential gene expression near sites of WT CRP occupancy only, CRP K52Q occupancy only, or occupancy of both transcription factors. Numbers of enrichment sites correlated with transcriptional activation or repression at Class I and Class II promoters (B) shared by WT CRP and CRP K52Q and (C) unique to CRP K52Q. (D) Impact of CRP K52Q binding sites within coding sequences (CDS) on transcription. A two-fold change in transcription with FDR < 0.05 was considered to be significant.

### The CRP K52Q substitution alters *V. cholerae* sugar utilization

CRP is best known for activating utilization of carbon sources other than glucose. The CRP K52Q substitution dramatically reshaped the transcription profile of sugar transport and utilization genes, suggesting a shift in carbon preferences. To illustrate this, we constructed a heat map comparing regulation of such genes by CRP (CRP/Δ*crp*) and the additional regulation imposed by a CRP K52Q substitution (CRP K52Q/WT CRP). As compared with the Δ*crp* mutant, WT CRP activated genes involved in transport and catabolism of citrate, maltose and galactose, which are transported independently of the PTS, as well as the PTS-dependent sugars mannitol, mannose, and cellobiose (Fig S4). In addition, CRP activated transcription of an alternative EllC glucose transporter, VC1821, that is reportedly independent of PTS Enzyme l, and nearby EllA and B homologs (VC1822 and VC1823) with unknown sugar specificity (5). In contrast, CRP K52Q principally activated transcription of genes involved in transport and utilization of PTS-independent carbon sources such as citrate, maltose, and galactose and repressed those involved in utilization of PTS-dependent carbon sources such as mannitol, mannose, glucose, and cellobiose. In addition, transcription of the fructose repressor, FruR, was increased, which would be predicted to decrease utilization of the PTS-dependent sugar fructose. Transcription of gene clusters involved in uptake and catabolism of only two PTS-dependent sugars, trehalose and the cell wall building block N-acetylmuramic acid, was activated by CRP K52Q. Both are implicated in the cellular response to osmotic stress (45, 46).

To test the functional implications of our observations, we measured *V. cholerae* growth on various sugars. We found that a CRP K52Q mutant grew more poorly than WT on minimal medium supplemented with fructose but better on that supplemented with trehalose (Fig S5). CRP K52Q grew better on several concentrations of maltose but only when inoculated from a pre-culture containing maltose. We conclude that the CRP K52Q substitution and, by extension, CRP K52Ac remodels sugar utilization by *V. cholerae*.

### The CRP K52Q substitution reverses repression of virulence gene expression by CRP but does not increase virulence in an infant mouse model

*V. cholerae* depends on the toxin co-regulated pilus (TCP) for colonization of the infant mouse and rabbit intestines (47). The VPS-dependent biofilm, whose role in attachment to abiotic surfaces has been intensively studied, is also conditionally required for colonization of the mammalian intestine (48–50). The toxins RTX and hemolysin are minor virulence factors (51). CRP represses expression of the TCP, biofilm, and RTX toxin genes in laboratory cultures and activates expression of a hemolysin gene. Nevertheless, a Δ*crp* mutant is defective in colonization of the infant rabbit intestine (23, 33, 52). This suggests that CRP may not repress virulence gene expression *in vivo.* To determine how the CRP K52Q point mutation impacted virulence gene expression, we compared transcript abundance in WT CRP and the K52Q mutant (Fig S6A). In spite of repression of the transcription regulators ToxR (VC0984) and TcpP (VC0826), which are activators of virulence gene transcription, the CRP K52Q substitution activated virulence gene expression, bringing transcription closer to that of a Δ*crp* mutant (53, 54). To determine how this might translate into virulence in the infant mouse model, we carried out competition experiments co-inoculating the 4XQ or K52Q mutant and WT *V. cholerae.* As shown in Fig S6B, the 4XQ mutant had a slight colonization defect, while colonization by the K52Q mutant was similar to that by WT *V. cholerae*. While colonization is a complex phenotype, these results are consistent with acetylation of CRP K52Q *in vivo*. This would allow *V. cholerae* to retain maximal virulence gene expression while optimizing carbon utilization of carbon sources available in the intestine.

### The CRP K52Q mutant blocks the acetate switch

Like *E. coli*, in the presence of an excess of carbon sources, *V. cholerae* excretes acetate as a byproduct of fermentation rather than channeling it into the tricarboxylic acid (TCA) cycle (55, 56). When these carbon sources are expended, *V. cholerae* initiates acetate consumption by upregulating the acetyl-CoA synthesis gene *acs1* in a process known as the acetate switch (55). In *V. cholerae*, the acetate switch is activated by the two-component system CrbRS and CRP (31, 56). We reasoned that CRP K52Ac might be more abundant during sugar fermentation and, therefore, that one role of both CRP K52Ac and CRP K52Q might be to modulate the acetate switch. To examine this hypothesis, we explored the transcription of switch-specific genes in our RNA-seq data set. In fact, the abundance of transcripts involved in acetate utilization such as *crbR* and the acetyl-CoA synthase gene *acs1* was much higher in WT *V. cholerae* as compared with the CRP K52Q mutant (Fig 4A). We confirmed these results by qRT-PCR (Fig 4B). To demonstrate that this pattern of gene regulation did, in fact, delay the acetate switch, we measured acetate in the supernatants of LB cultures (Fig 4C). Acetate consumption was delayed in the CRP K52Q mutant as compared with WT *V. cholerae*. Growth on acetate involves synthesis of acetyl-CoA, funneling of acetyl-CoA into the TCA cycle, and upregulation of the glyoxylate shunt to avoid the carbon wasting steps in the TCA cycle (57). The CRP K52Q mutant was unable to use acetate as a sole carbon source for growth (Fig 4D). Taken together, these results are consistent with a model in which acetylation of CRP K52 functions to block the acetate switch.

**Figure 4:**
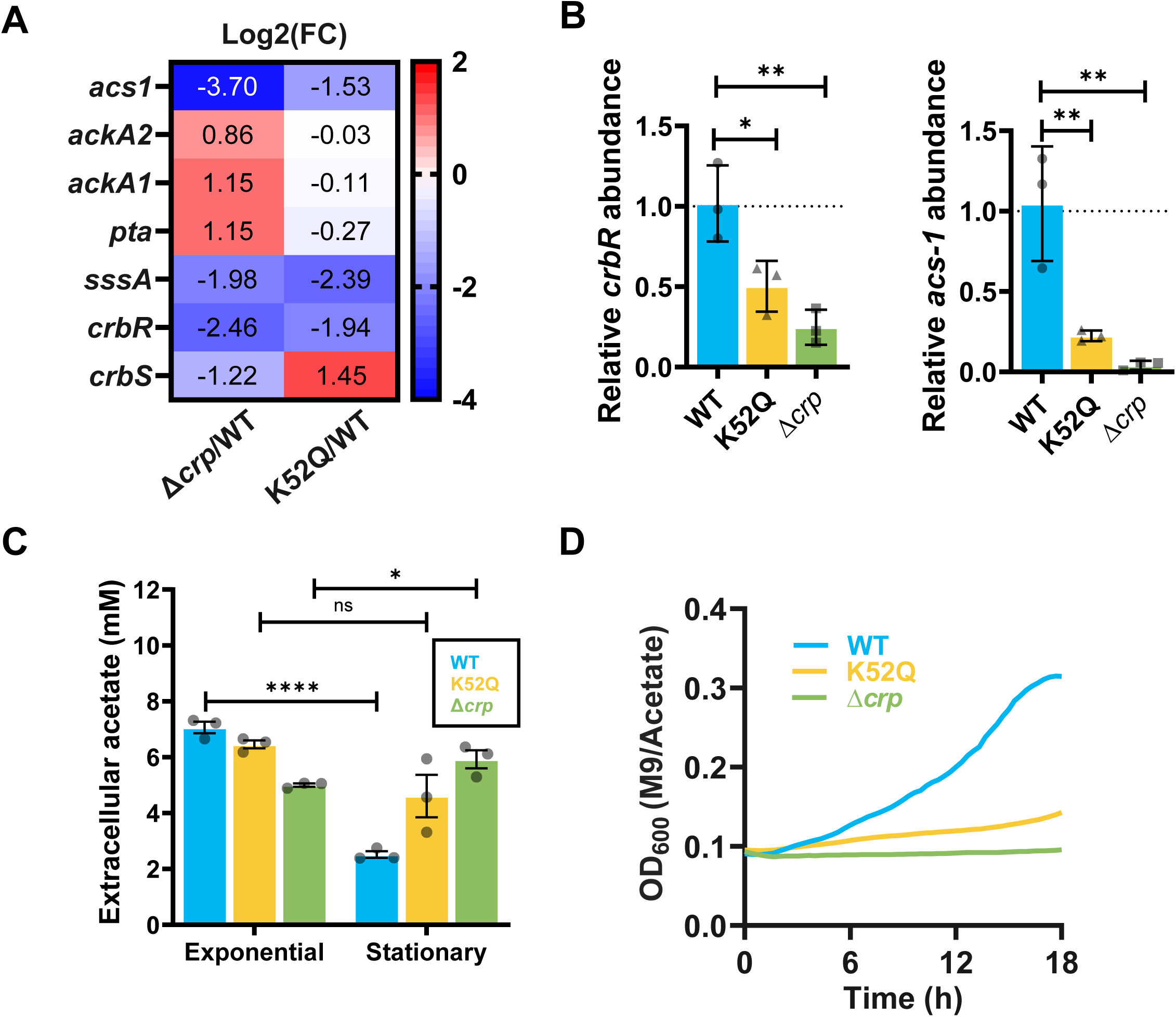
CRP K52Q delays the acetate switch. (A) Heat map showing differential regulation of genes essential for the acetate switch. Data is the mean of RNA-seq biological triplicates. (B) qRT-PCR analysis of *crbR* and *acs1* transcription in the indicated *V. cholerae* strains. The mean of biological triplicates is shown. An ordinary one-way ANOVA was used to determine significance. (C) Acetate concentrations in the supernatants of exponential and stationary phase cultures of WT *V.* cholerae and CRP K52Q, and Δ*crp* mutants. The mean of biological triplicates is shown. A student’s t test was used to determine significance. (D) Growth over time of the indicated strains in M9 medium supplemented with 0.4 % acetate. The average of biological triplicates is shown. **** p<0.0001, *** p<0.001, ** p<0.01, * p<0.05, ns not significant.

### A medium rich in fermentable sugars increases lysine acetylation but not of K52

Because a CRP K52Q mutant delays the acetate switch, we hypothesized that we might detect CRP K52 acetylation in a medium rich in a fermentable sugar. To test this, we purified CRP-V5 from exponential and stationary phase cultures in minimal medium supplemented with 0.4% sucrose and sent the purified protein for proteomic analysis. As shown in Figure S7, acetylation of K35, K89, and K188 and succinylation of K188 was detected in exponential phase. In stationary phase when the sucrose is depleted, the acetylation sites were no longer observed. However, acetylation of CRP K52 was not observed in any of these cultures. This demonstrates that while fermentable sugars in the environment are correlated with *V. cholerae* CRP lysine acetylation, acetylation of K52 is not detectable in the setting of sucrose abundance. We hypothesize that acetylation of CRP K52 demonstrates sugar specificity.

### CRP K52Q activates transcription of the novel sRNA gene *crbZ*, which is transcribed within and antisense to *crbS*

We noted that several binding sites unique to CRP K52Q were close to TSSs of novel transcripts. Six of these, shown in Fig S8, were positioned either within an intragenic region or within a CDS but running antisense to the CDS. The distance of all the CRP K52Q binding sites from a TSS was most consistent with that of a Class ll promoter. This is supports our previous observations that the CRP K52Q mutant activates transcription from Class ll promoters, possibly by engaging AR3.

We further explored a putative sRNA encoded anti-sense to *crbS* (Fig 5A). This gene was designated *crbZ* to reflect its location. We first confirmed its transcript abundance by qRT-PCR using primers designed to amplify the *crbS* region containing *crbZ* and primers designed to amplify a region of *crbS* downstream of *crbZ.* As shown in Fig 5B, when primers spanning *crbZ* were used to measure *crbZ*+*crbS* transcript abundance, that measured for the CRP K52Q mutant was approximately 13-fold higher than that for WT *V. cholerae* or a Δ*crp* mutant. In contrast, when primers in the histidine kinase domain (HK) outside *crbZ* were used, there was no significant difference in transcript abundance between the CRP K52Q mutant and WT *V. cholerae* or a Δ*crp* mutant. This supports the presence of a sRNA in the N-terminus of *crbS* that is regulated by CRP K52Q. To further establish the presence of this sRNA, we performed a ligation to circularize CrbZ, generated cDNA, and amplified the region spanning the ligated ends using outward facing primers at the 5 and 3 ends of *crbZ* (Fig 5C). As shown in Fig 5D, the primers amplified a product. To prove that this product was ligated CrbZ, it was sequenced. The sequence of the amplification product matched the ligated ends of CrbZ (Fig S9). This provides strong evidence that *crbZ* is transcribed.

**Figure 5:**
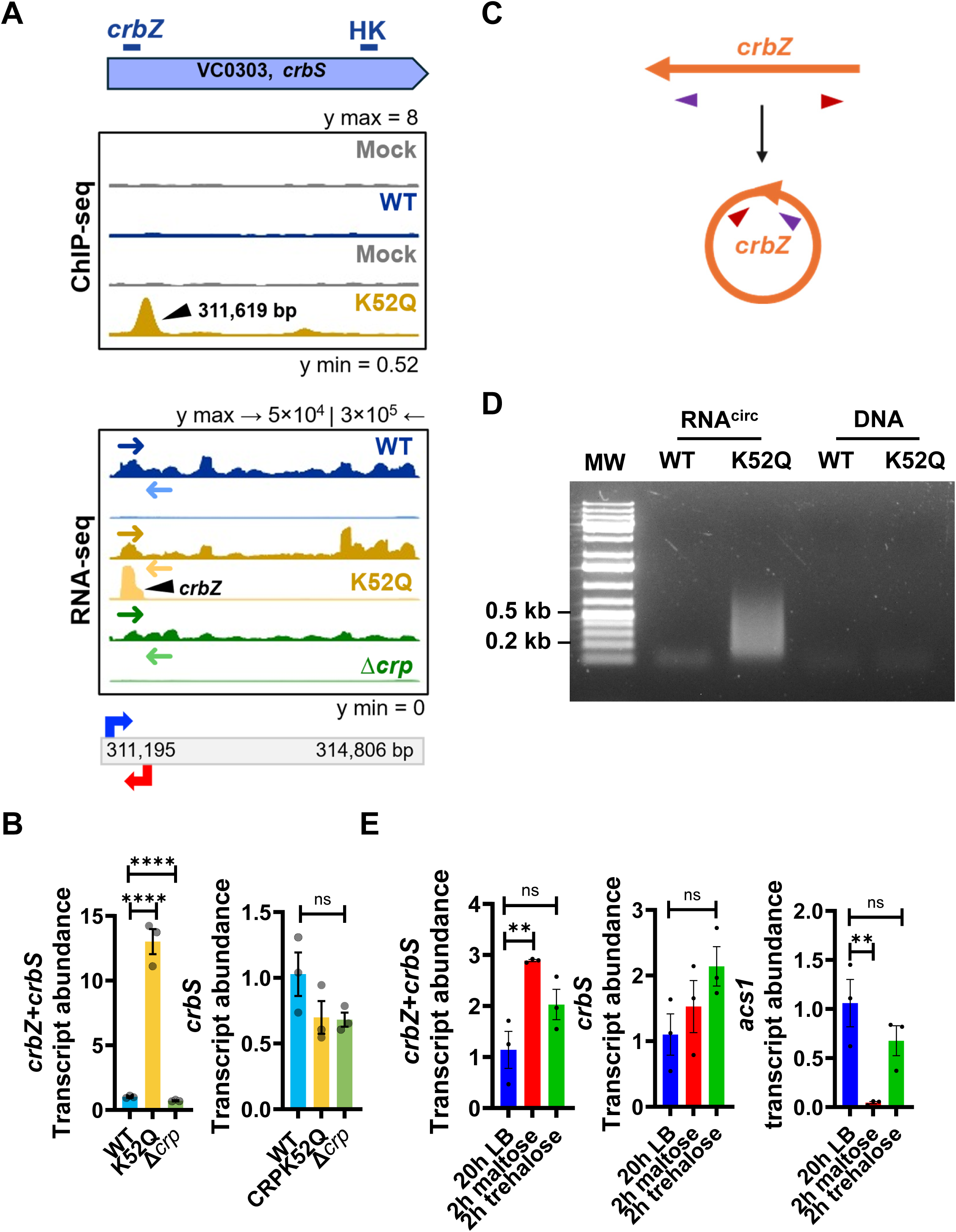
The small RNA CrbZ is encoded antisense and within CrbS and activated by CRP acetylation. (A) ChIP-seq and RNA-seq traces showing enrichment of CRP K52Q at a class ll promoter and transcription of the *crbZ* sRNA gene, respectively. Traces are representative of measurements in biological triplicate. These data are also shown in Fig S 8A. (B) qRT-PCR quantification of transcription of *crbZ+crbS* and *crbS* in the indicated strains. The mean of biological triplicates is shown. Error bars represent the mean. A one-way ANOVA or student’s t test was used to assess significance. (C) Schematic showing the experimental design and (D) an agarose gel to document the presence of the *crbZ* sRNA by ligation and PCR amplification. Arrows indicate the position of the PCR primers. (E) qRT-PCR quantification of *crbZ + crbS, crbS*, or *acs1* transcription in WT *V. cholerae* cultured for 20h in LB broth or in M9 minimal medium supplemented with 0.4% maltose or trehalose for 2 h. The mean of biological triplicates is shown. Error bars represent the mean. A one-way ANOVA was used to assess significance. **** p<0.0001, *** p <0.001, ** p<0.01, * p< 0.05, ns not significant.

### Like CRP K52Q, CrbZ inhibits the acetate switch and improves growth on maltose

Based on its position within the *crbS* gene, we hypothesized that CrbZ might regulate acetate uptake. To test this, we cloned *crbZ* into an IPTG-inducible vector. We first confirmed overexpression of *crbZ* from the plasmid (Fig S10A). We then tested its impact on *crbS, crbR,* and *acs1* expression and acetate uptake. *crbZ* overexpression decreased both *acs1* transcript abundance and delayed acetate uptake but did not affect expression of *crbRS* (Fig S10A and B). Similar to the CRP K52Q mutant, *crbZ* overexpression improved growth in maltose (Fig 10C). Taken together, these data suggests that CrbZ plays a role in repression of acetate uptake by the CRP K52Q mutant.

### *crbZ* is expressed in minimal medium containing maltose

Our attention was drawn to *crbZ* precisely because it was transcribed when the CRP K52Q mutant but not WT *V. cholerae* was cultured in LB broth. We hypothesized that *crbZ* might be transcribed by WT *V. cholerae* in minimal medium supplemented with maltose, a carbon source that promoted growth of the CRP K52Q mutant. In fact, we found that in exponential phase cultures of WT *V. cholerae* in minimal medium supplemented with maltose but not trehalose, the region of *crbS* that includes *crbZ* was expressed approximately 3-fold more highly than in the LB overnight culture that was used to inoculate the minimal medium cultures (Fig 5E). In contrast, the distal region of *crbS* was not differentially transcribed for any of these conditions. To measure the transcript abundance of *crbZ* and *crbS* separately, we performed strand-specific qRT-PCR. To validate the technique, we first measured *crbZ* and *crbS* transcript abundance in WT *V. cholerae* and the CRP K52Q mutant. In WT *V. cholerae* the ratio of *crbZ*/*crbS* was approximately 0.2, while in the CRP K52Q mutant it was approximately 14 (Fig S11A). This is consistent with our RNA-seq results. In minimal medium supplemented with 0.8% maltose, the ratio of *crbZ*/*crbS* was approximately 14 in exponential phase cultures and approximately 6 in stationary phase cultures (Fig S11B). This confirms upregulation of *crbZ* when *V. cholerae* is cultured in the presence of maltose.

Furthermore, *acs1* was downregulated only in the minimal medium culture supplemented with maltose but not trehalose (Fig 5E). Taken together, our results suggest that *crbZ* is expressed in a sugar-specific manner and represses the acetate switch.

## Discussion

CRP is one of the best studied bacterial transcription regulators. It activates transcription at Class l and Class ll promoters through ARs 1 and 2, which have been defined by structure-function studies (16, 37, 58). Approximately 25 years ago, investigators observed that substitution of CRP K52 for a neutral residue resulted in transcription activation specifically at Class ll promoters when CRP contained inactive ARs 1 and 2. This led to the discovery of AR 3, which was dismissed as dispensable for transcription activation by native CRP (37). Here we show that a CRP K52Q substitution, which mimics acetylation of K52, results in hundreds of new chromosomal binding sites. Approximately half of these binding sites are in promoters within intergenic regions, while the other half are within genes. Almost all of the novel binding sites that impact transcription are located proximal to TSSs in positions consistent with class ll promoters. Furthermore, the occupancy of CRP K52Q at many of these sites positively regulates the production of RNAs that have not previously been reported. Our results suggest that acetylation of CRP K52 engages AR3 leading to transcription of novel RNAs that rewire sugar utilization, block the acetate switch, and increase virulence gene expression. We further show that at least one sRNA activated by CRP K52Q, *crbZ*, is expressed by WT *V. cholerae* in maltose-containing medium and represses the acetate switch. All of these findings are consistent with a role for CRP K52Ac in remodeling *V. cholerae* carbon utilization and virulence to optimize survival in the human intestine.

In LB broth cultures, CRP K52Q occupancy was noted at hundreds of sites not occupied by WT CRP. Using MEME analysis to examine the consensus binding sequences for WT CRP and CRP K52Q, we discovered that many CRP K52Q binding sites belong to one of two consensus sequences, one that is similar to that of WT CRP and an additional one in which half of the inverted repeat is conserved and the other half is highly degenerate. Initial studies describing activation of transcription by AR3 documented an interaction with region 4 of the σ^70^ subunit of RNAP at Class ll promoters (18, 59). However, not all the binding sites unique to the CRP K52Q mutant are located in Class ll promoters near TSSs (38). We examined the possibility that the interaction of the downstream CRP monomer with the σ^70^ subunit of RNAP positioned either at a promoter or within a gene in a paused elongation complex might stabilize binding of CRP K52Q to DNA. Our results suggest this is not the case. Instead, CRP K52Q appears to be able to bind DNA when only one half of the consensus inverted repeat is a good match regardless of an interaction with σ^70^. Furthermore, it is able to modulate transcript abundance when bound at these sites if they are positioned within promoters demonstrating that CRP K52Q is competent to activate transcription at these sites. Further studies are required to understand the mechanism by which the K52Q substitution promotes binding and activates transcription at degenerate operator sites.

The CRP K52Q substitution, which mimics acetylation, modifies the *V. cholerae* transcriptome by altering expression of many genes known to be controlled by WT CRP and generating several putative novel RNAs. One of these sRNAs, CrbZ, is responsible for a subset of the changes observed in the CRP K52Q transcriptome. Because so many of CRP K52Q’s binding sites are within genes and activate antisense transcription, it is interesting to speculate that sRNA regulation plays a dominant role in modulation of the CRP regulon by K52Ac. By using conditionally expressed sRNAs, CRP can retain WT expression of most of its regulon, while modifying transcription of only those genes that are required to respond to a change in carbon source availability.

Bacterial carbon overflow metabolism is defined by fermentation of sugars during rapid growth in oxygen-rich environments (60). This results in generation of large amounts of short chain fatty acids such as acetate and is predicted to increase protein acetylation in general. We show here that CRP acetylation is increased during exponential growth in defined medium supplemented with sucrose. Because we did not detect acetylation of CRP K52 under these conditions, we hypothesize that the abundance of *V. cholerae* CRP K52Ac depends on the availability of specific sugars.

When carbon sources become scarce, uptake of acetate is initiated in a process known as the acetate switch (55). We predict that acetylation of CRP signals excess carbon availability to enhance carbon overflow metabolism while suppressing the acetate switch. In fact, the response of a K52Q mutant is concordant with these predictions. Interestingly, while the transcript abundance of genes involved in uptake and utilization of many PTS-dependent sugars is decreased, the abundance of those involved in uptake of trehalose is increased. This is intriguing because trehalose has been shown to counteract the detrimental effects of protein acetylation (61). Therefore, a selective increase in trehalose uptake and generation may be beneficial in the setting of excess carbon availability. Taken together, these data suggest that CRP K52Ac prepares *V. cholerae* to thrive in specific carbon-rich environments.

CRP represses virulence gene expression and yet, in spite of increased virulence gene expression, a Δ*crp* mutant has a massive colonization defect *in* vivo (23, 33, 52). Furthermore, transposon insertions in maltose metabolism genes, whose transcript abundance is increased in the CRP K52Q mutant, had a competitive disadvantage in colonization of the infant rabbit intestine (33). This suggests that virulence factors may not be repressed by CRP *in vivo* and that regulation of sugar utilization by CRP is essential for *V. cholerae* growth in the mammalian intestine. Carbon sources such as citrate, galactose, and maltose, which may promote CRP K52 acetylation, are abundant in the human diet and principally absorbed in the small intestine where *V. cholerae* TCP-dependent colonization is observed (62). Acetylation of CRP *in vivo* would allow for maximal virulence gene expression while preserving regulation of sugar utilization by CRP. *V. cholerae* is found in human and environmental host intestines but is also cultured from the aquatic environment during epidemics. As carbon is limiting in most natural aquatic environments, it seems more likely that activation of carbon overflow metabolism by acetylated transcription factors such as CRP is operative in the host intestine. Therefore, we propose that acetylation of CRP K52 *in vivo* activates a transcriptional pattern that aids both colonization of and replication in the mammalian intestine.

## Materials and Methods

A complete reference list of strains used in this study is provided in Table S5. *Vibrio cholerae* derived from the O1 biovar El Tor strain C6706str2 was cultured in Lysogeny Broth (LB; Difco 244620; 10 g tryptone, 5 g yeast extract, 10 g NaCl per liter) or in M9 minimum medium (1× M9 salts; Difco) supplemented with 1 mM MgSO_4_, 0.1 mM CaC_2_ and 0.4% [w/v] of the indicated carbon source. *Escherichia coli* strains used for cloning were grown in LB only. Cultures were incubated at 27 °C (*V. cholerae*) or 37 °C (*E. coli*) with shaking at 200 rpm. *V. cholerae* strains were maintained in the presence of streptomycin (100 µg/mL), and, when required, carbenicillin (100 µg/mL) was added for plasmid selection. Biological triplicates were performed for all experiments.

### Plasmid construction and mutant generation

PCR-amplified fragments were cloned into the *Xho*I site in the pWM91 vector to generate constructs for in-frame deletions or chromosomal complementation using New England Biolabs (NEB) HiFi DNA Assembly (NEB, E2621L). For gene overexpression, PCR products containing the gene and its native promoter were cloned into the *Xho*I site of the pFLAG-CTC IPTG-inducible vector. *V. cholerae* strains were generated via homologous recombination using the Sanger-sequenced plasmids introduced by conjugation from *E. coli* SM10 λpir donor strains. Selection with 12% sucrose and PCR screening were used to confirm successful in-frame deletion. Chromosomal complementation was achieved by allelic exchange at the native locus in a deletion background strain.

### Chromatin immunoprecipitation and sequencing (ChIP-seq)

ChIP was performed as previously described with minor modifications (34). Briefly, protein-DNA complexes were cross-linked and immunoprecipitated with anti-V5 antibodies. The cross-links were subsequently reversed and co-purifying DNA was purified using the Qiaquick PCR Purification kit. DNA libraries were prepared with the NEBNext Ultra II DNA Library Prep Kit (NEB E7645S) according to the manufacturer’s instructions and sequenced on an Illumina NovaSeq 6000 V1.5 SP 100 cycles Full Flow Cell (650 million reads) at the Harvard Biopolymers Facility.

### ChIP-seq computational analysis

Paired-end reads were aligned to the *V. cholerae* reference genome (NC_002505.1; NC_002506.1) using Bowtie2 (v2.4.1). Following mapping, an in silico size selection was performed to retain fragments shorter than 400 nt in length. The 3 mock biological replicates were merged and subsampled (samtools v1.10, using htslib 1.10.2), so that they had approximately 2 million reads more than any single ChIP replicate file. Peak calling was performed with QuEST (version 2.424) to identify putative DNA-binding sites defined as regions exhibiting more than two-fold enrichment relative to background from an untagged mock sample. Peaks assignments were performed by finding the closest gene to the point of maximal enrichment using custom perl scripts and/or manual curation of the data. Peaks positioned such that no genes were located within 1000 bp of the site of maximal enrichment were excluded. For each strain, peaks with enrichment factor (EF) > 4 and reproducible in at least two of three biological replicates were retained for downstream analyses. All custom Python scripts used for data processing, analysis, and figure generation are available at https://github.com/renatoerss/vch-crp-acetylation, and summarized below.

Peaks identified in each strain were paired by genomic overlap based on the reported region start and end coordinates. Shared and strain-specific datasets were summarized with Upset plots or Venn diagrams. Transcription start sites (TSSs) were obtained from the differential RNA-seq dataset of *V. cholerae* and used to compute TSS distances and define promoter classes (Class I and II) (38). Genomic categories were defined as promoter (-200 to -1 nt relative of TSS), 5’-UTR (+1 from TSS to the base preceding the annotated start codon), coding sequence (CDS), or other, if not within these regions.

For motif discovery, 100-bp windows centered on the ChIP-seq peak maximum position were extracted and analyzed with MEME-ChIP (MEME suite, https://meme-suite.org/meme/), using MEME (anr model, 16-22 bp motif width), Centrimo for positional enrichment, and Tomtom for motif comparison against curated bacterial motif databases (63, 64). Integration with RNA-seq used per-gene log_2_ FC (K52Q/WT). Peak categories (WT-only, WT + K52Q, and K52Q-only) are represented in the figure.

### RNA isolation and transcriptome analysis

Total RNA from mid-exponential-phase (OD_600_ 0.4-0.6) culture was isolated using the Direct-zol RNA Miniprep Plus kit (Zymo Research) following the manufacturer’s instructions. Genomic DNA was removed by in-column DNase treatment, and RNA quality was assessed using a TapeStation system (Agilent Technologies).

RNA-seq libraries were prepared using a modified version of the RNAtag-Seq protocol starting from 500 ng of total RNA (65). Briefly, RNAs were fragmented, dephosphorylated, and ligated to DNA adaptors carrying 5’-AN₈-3’ barcodes with a 5’ phosphate and a 3’ blocking group. Barcoded samples were pooled and subjected to rRNA depletion using the Ribo-Zero rRNA Removal Kit (Epicentre).

Sequencing was performed on an Illumina NextSeq 500 platform, generating paired-end reads. The average sequencing depth was ∼12 million fragments/sample, with three biological replicates per condition. Reads were aligned to the *Vibrio cholerae* O1 biovar El Tor str. N16961 (accession numbers NC_002505.1 and NC_002506.1) genome using BWA, and read counts were assigned to annotated genomic features using custom scripts provided by the Microbial ‘Omics Core (MOC) at the Broad Institute (66). Differential expression analysis was performed using DESeq2, and expression values were reported as fragments per kilobase per million mapped reads (FPKM)(67). Visualization of RNA-seq coverage traces and manual gene-level inspection was performed using the Integrative Genomics Viewer (68).

To assess sample reproducibility and global transcriptional differences within the CRP regulon, a principal component analysis (PCA) was performed using genes with greater than two fold change and adjusted *P-*value < 0.05. The analysis was conducted on normalized, log-transformed read counts. The first two principal components accounted for the majority of the variance and separated biological replicates by strain. PCA was computed in R (version 4.4.3) using stats::prcomp.

### Quantitative reverse transcription PCR (qRT-PCR)

Total RNA was extracted as previously described. 500 ng of total RNA was reverse-transcribed using the SuperScript III First-Strand Synthesis System (Invitrogen). qPCR reactions were performed in 10 µL volumes using PowerUp SYBR Green Master Mix for qPCR (Applied Biosystems) on a QuantStudio 5 Real-Time PCR System (Applied Biosystems). Gene-specific and housekeeping primers are listed in Table S5. Relative transcript abundance was calculated by the ΔΔCt method, normalized to *clpX* (VC1921). All reactions were performed in technical duplicates using three independent biological samples.

### Extracellular acetate quantification

Extracellular acetate levels were determined using EnzyChrom Acetate Assay Kit (BioAssay Systems, EOAC-100) following manufacturer’s protocol. Culture supernatants were collected from mid-exponential (OD_600_ 0.4-0.6) or stationary phase cultures (18 h incubation under standard conditions). Samples were clarified by centrifugation and filtered through 0.22 µm PVDF membranes. Acetate concentration was determined from a sodium acetate standard curve and absorbance was measured at 570 nm using a SpectraMax iD5 microplate reader (Molecular Devices).

### Circularization RT-PCR (cRT-PCR)

The 5’ and 3’ ends of the *crbZ* transcript were mapped according to a previously described RNA circularization protocol (69). Briefly, DNAse I treated total RNA (2 µg) was decapped with 2 U of RNA 5’-pyrophosphohydrolase (RppH; New England Biolabs) in 1× NEBuffer 2 for 1 h at 37 °C to generate 5’-monophosphorylated ends. The RNA was purified using the RNA Clean & Concentrator kit (Zymo Research) and circularized overnight at 16 °C using 1 U of T4 RNA Ligase 1 (NEB) in 1× RNA Ligase Buffer supplemented with 1 mM ATP, 8 µL PEG 8000 (50%), 2 µL DMSO, and 0.75 U RNase inhibitor in a total volume of 35 µL. Circularized RNA was cleaned again and eluted in nuclease-free water. cDNA synthesis was carried out with 500 ng of circular RNA using the SuperScript III First-Strand Synthesis System (Invitrogen) and random hexamer primers, following manufacturer’s protocol. The resulting cDNA was amplified by PCR using outward-facing primers specific for *crbZ* and designed to flank the ligated junction (Supplementary Table S4). PCR reactions were carried out with Q5 High Fidelity DNA Polymerase (New England Biolabs) for 30 cycles, and products were resolved on 2% agarose gels. Control PCR reactions with genomic DNA were included to confirm primer specificity. Specific amplicons (150-400 bp) detected in the CRP K52Q sample were purified and sent for DNA sequencing.

### Statistical analysis

All experiments included a minimum of biological triplicates. Graphs were constructed and statistical analysis was carried out using GraphPad Prism (version 10.6.1). The appropriate statistical test was applied to evaluate significance as noted in the figure legends.

## Data availability statement

ChIP-seq and RNA-seq data have been deposited in NCBI’s Gene Expression Omnibus and are accessible through GEO Series accession number GSE310654.

## Acknowledgements

This work was supported by NIH R01 AI112652 to P.I.W. and NIH R01AI018045 to J.J.M.

## Author Contributions

RS, JG, and PIW designed the research. SLD and JJM provided experimental advice and resources. RS, JG, CA, PB, MJG, and WPR performed the research. RS, MJG, PB, and SLD contributed new analytic tools. RS, JG, MJG, PB, WPR, SD, and PIW analyzed the data. RS and PIW wrote the manuscript. All authors reviewed the manuscript.

## Declaration of interests

The authors declare no competing interests.

---

**Figure S1: A CRP K52Q point mutant remains membrane-associated.**

**Figure S2. A CRP C-terminal V5 tag has only a small impact on the *V. cholerae* transcriptome.**

**Figure S3: Chromosomally-encoded CRP acetylation mutants are expressed at comparable levels, grow at comparable rates in LB broth, and do not affect CRP acetylation at other sites.**

**Figure S4: The CRP K52Q mutant modulates the transcript abundance of genes required for utilization of fructose, trehalose, citrate, maltose mannitol, mannose, and cellobiose.**

**Figure S5: The CRP K52Q substitution alters growth of *V. cholerae* in a variety of sugars.**

**Figure S6: The CRP K52Q mutant increases the transcript abundance of genes required for synthesis of the toxin co-regulated pilus, the VPS-dependent biofilm, and the RTX toxin.**

**Figure S7: In minimal medium supplemented with sucrose, *V. cholerae* CRP acetylation decreases in stationary phase.**

**Figure S8: Six putative RNAs are uniquely activated by CRP K52Q. Figure S9: Sequence alignment of the circularized RNA junction.**

**Figure S10: The sRNA CrbZ regulates the acetate switch and maltose metabolism.**

**Figure S11: Strand-specific RT-qPCR demonstrates activation of *crbZ* transcript abundance in maltose-containing medium.**

**Table S1: WT CRP, CRP 4XR, and CRP 4XQ ChIP-seq analysis.**

**Table S2: Deseq comparisons of WT (C6706), WT CRP-V5, CRP K52Q-V5, CRP 3XQ-V5, CRP 4XQ-V5, CRP 4XR-V5 and** Δ***crp* transcriptomes in exponential LB broth cultures.**

**Table S3: WT and K52Q ChIP-seq data**

**Table S4: List of meme sites for WT CRP and CRP K52Q Table S5: Strains, plasmids, and primers used in this study**

## Notes

https://www.ncbi.nlm.nih.gov/geo/query/acc.cgi?acc=GSM9305471

### Competing Interest Statement

The authors have declared no competing interest.

