## Supplemental Figures for "Actuation of CRP activating region 3 by acetylation modulates *V. cholerae* sugar utilization and virulence"

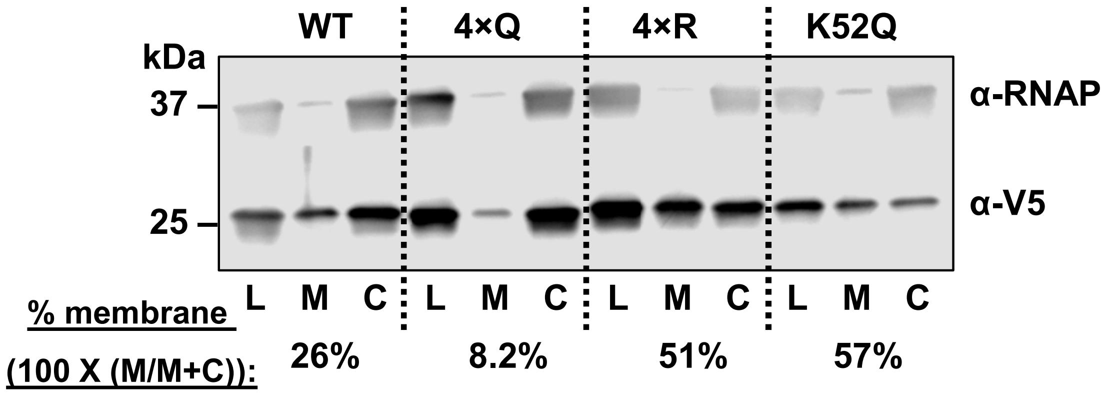


**Figure S1: A CRP K52Q point mutant remains membrane-associated.** Western blot analysis of total lysate (L), membrane (M), and cytoplasmic (C) fractions from *V. cholerae* strains expressing chromosomally encoded V5-tagged CRP alleles including WT CRP (WT), CRP with K22, K26, K35, and K52 mutated to Q (4XQ) or R (4XR), or just K52 mutated to Q (K52Q). Cells were cultured in LB broth and harvested in mid-exponential phase. Tagged CRP was detected using an anti-V5 antibody (~28 kDa), and the RNA polymerase α-subunit (37 kDa) served as a control for fractionation. A representative blot from at least biological triplicates is shown. The intensity of bands representing the membrane-associated (M) and cytoplasmic (C) CRP fractions was quantified using FIJI (https://imagej.net/software/fiji/), and membrane association (%) was calculated as M/(M+C) $\times$ 100.

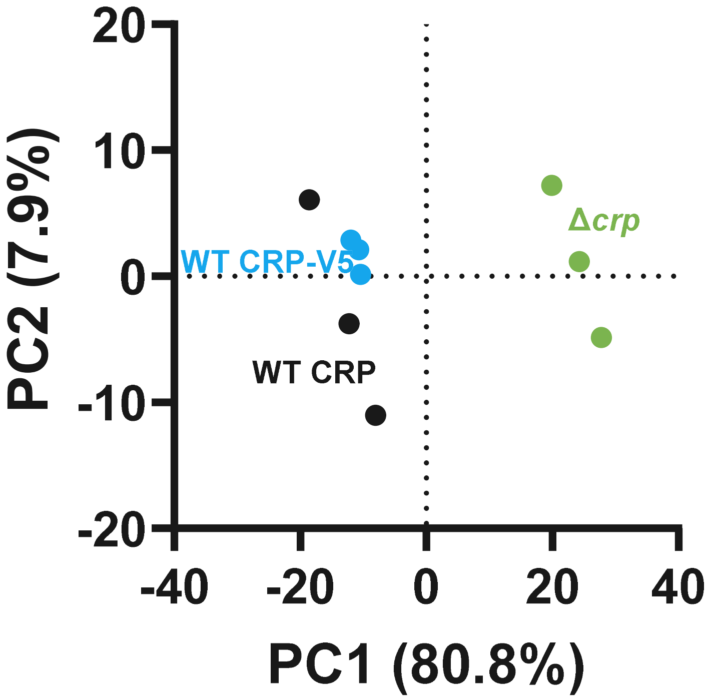


**Figure S2. A CRP C-terminal V5 tag has only a small impact on the *V. cholerae* transcriptome.** Principal component analysis (PCA) illustrating the differences between the transcriptomes of *V. cholerae* encoding untagged CRP, CRP-V5, or an in-frame deletion of CRP (Δ*crp*). Biological triplicates were included in RNA-seq analysis. The WT CRP-V5 and Δ*crp* data were also used to generate Fig 1D.

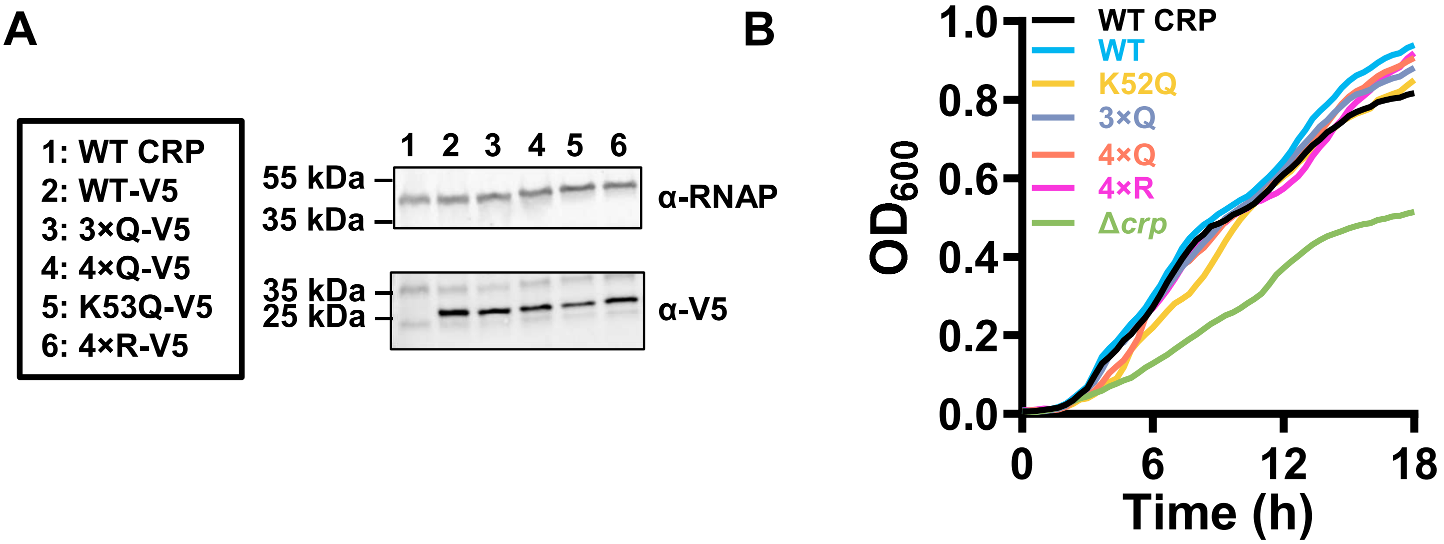


C

| Strain | Peptide identified | Acetylation site |
|---|---|---|
| CRP-V5 | R.ILK#M*LEEQNLISAHGK.T | K188 |
| | R.AK#TPCEVAEISFK.K | K89 |
| CRP 3×Q-V5 | R.ILK#M*LEEQNLISAHGK.T | K188 |
| | R.AK#TPCEVAEISFK.K | K89 |
| CRP 4×Q-V5 | R.ILK#M*LEEQNLISAHGK.T | K188 |
| | R.AK#TPCEVAEISFK.K | K89 |
| CRP 4×R-V5 | R.ILK#M*LEEQNLISAHGK.T | K188 |
| | R.AK#TPCEVAEISFK.K | K89 |
| CRP K52Q-V5 | R.ILK#M*LEEQNLISAHGK.T | K188 |
| | R.AK#TPCEVAEISFK.K | K89 |

**Figure S3: Chromosomally-encoded CRP acetylation mutants are expressed at comparable levels, grow at comparable rates in LB broth, and do not affect CRP acetylation at other sites.** (A) Immunoblot of mid-exponential LB cultures of *V. cholerae* expressing chromosomally-encoded CRP, CRP-V5, CRP with K22, K26, and K35 mutated to Q (3XQ), CRP with K22, K26, K35, and K52 mutated to Q (4XQ), K52 mutated to Q (K52Q), and (v) K22, K26, K35, and K52 mutated to R (4XR). The blot was probed with antibodies against the V5 tag and α-RNAP. (B) LB Growth curves of the strains listed in (A) as well as a Δ*crp* mutant. The mean of biological triplicates is shown. (C) Proteomic analysis of post-transcriptional modifications found in WT *V. cholerae* and the indicated point mutants. CRP was purified from mid-log cultures.

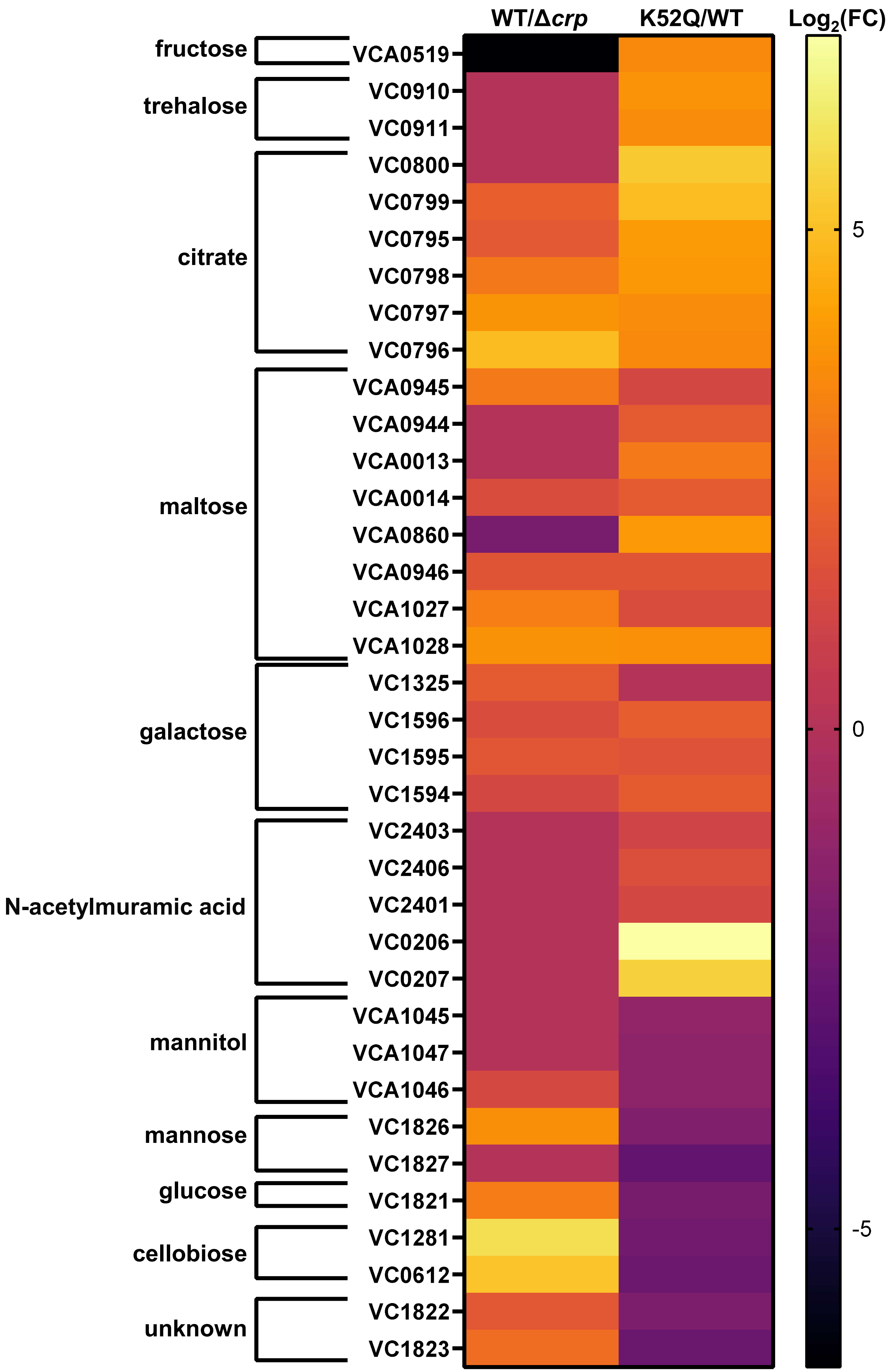

WT/Δcrp
K52Q/WT
Log2(FC)
fructose
trehalose
citrate
maltose
galactose
N-acetylmuramic acid
mannitol
mannose
glucose
cellobiose
unknown
VCA0519
VC0910
VC0911
VC0800
VC0799
VC0795
VC0798
VC0797
VC0796
VCA0945
VCA0944
VCA0013
VCA0014
VCA0860
VCA0946
VCA1027
VCA1028
VC1325
VC1596
VC1595
VC1594
VC2403
VC2406
VC2401
VC0206
VC0207
VCA1045
VCA1047
VCA1046
VC1826
VC1827
VC1821
VC1281
VC0612
VC1822
VC1823
5
0
-5

**Figure S4: The CRP K52Q mutant modulates the transcript abundance of genes required for utilization of many carbon sources.** Heat map illustrating the relative transcript abundance of the indicated genes for WT CRP vs a Δ*crp* mutant and a CRP K52Q mutant vs WT CRP. Colors reflect the $log_2FC$. Where no significant difference was noted, $log_2FC=0$ was used. EdgeR was used to identify significantly differently regulated genes

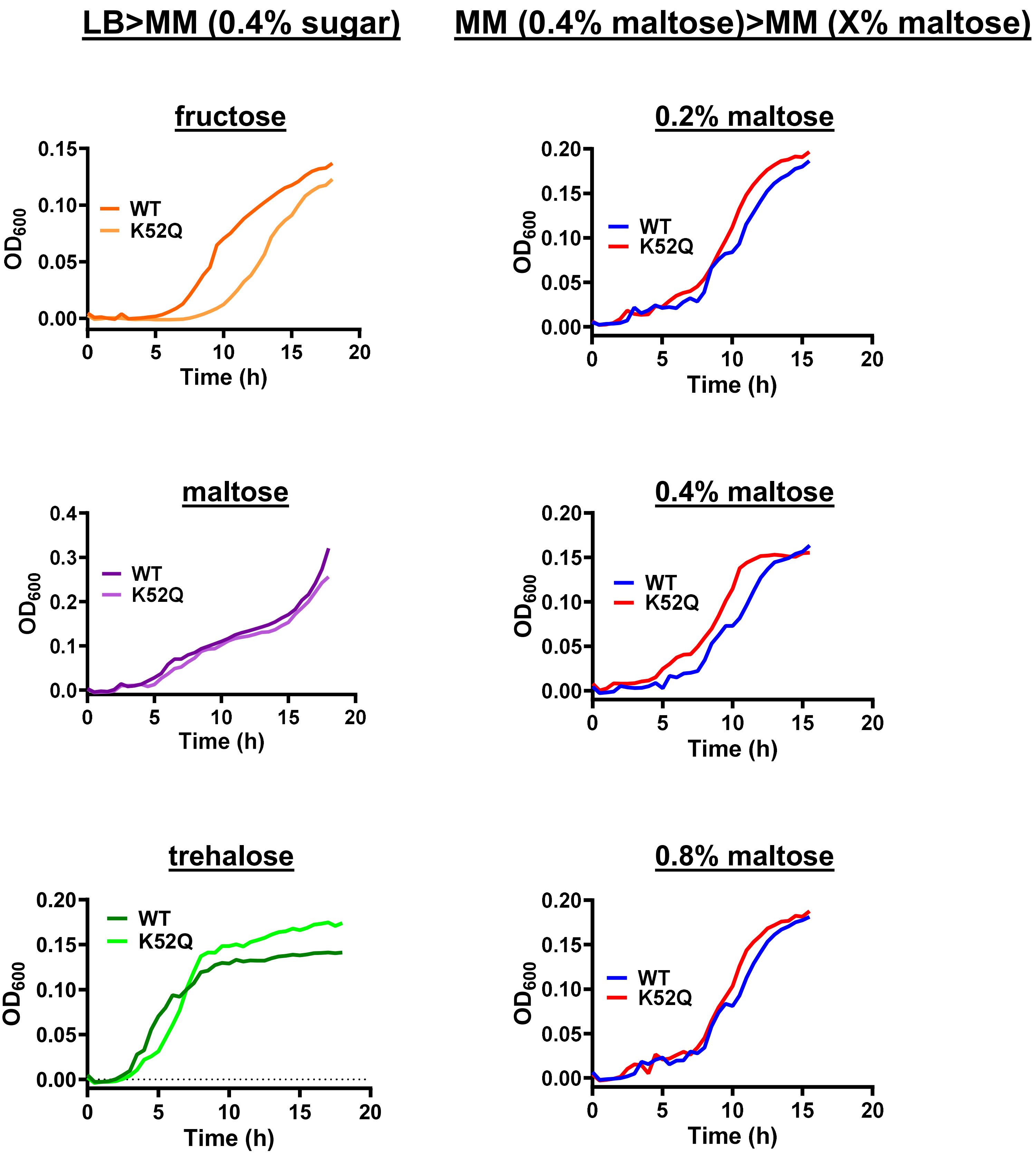


**Figure S5: A CRP K52Q substitution alters growth of *V. cholerae* in a variety of sugars.** Growth of WT *V. cholerae* and the K52Q mutant in M9 medium containing the indicated sugar. All growth curves represent averages of biological triplicates. LB>MM indicates back-dilution from an LB pre-culture, while MM>MM indicates back-dilution from a preculture in M9 minimal medium supplemented with 0.4% maltose.

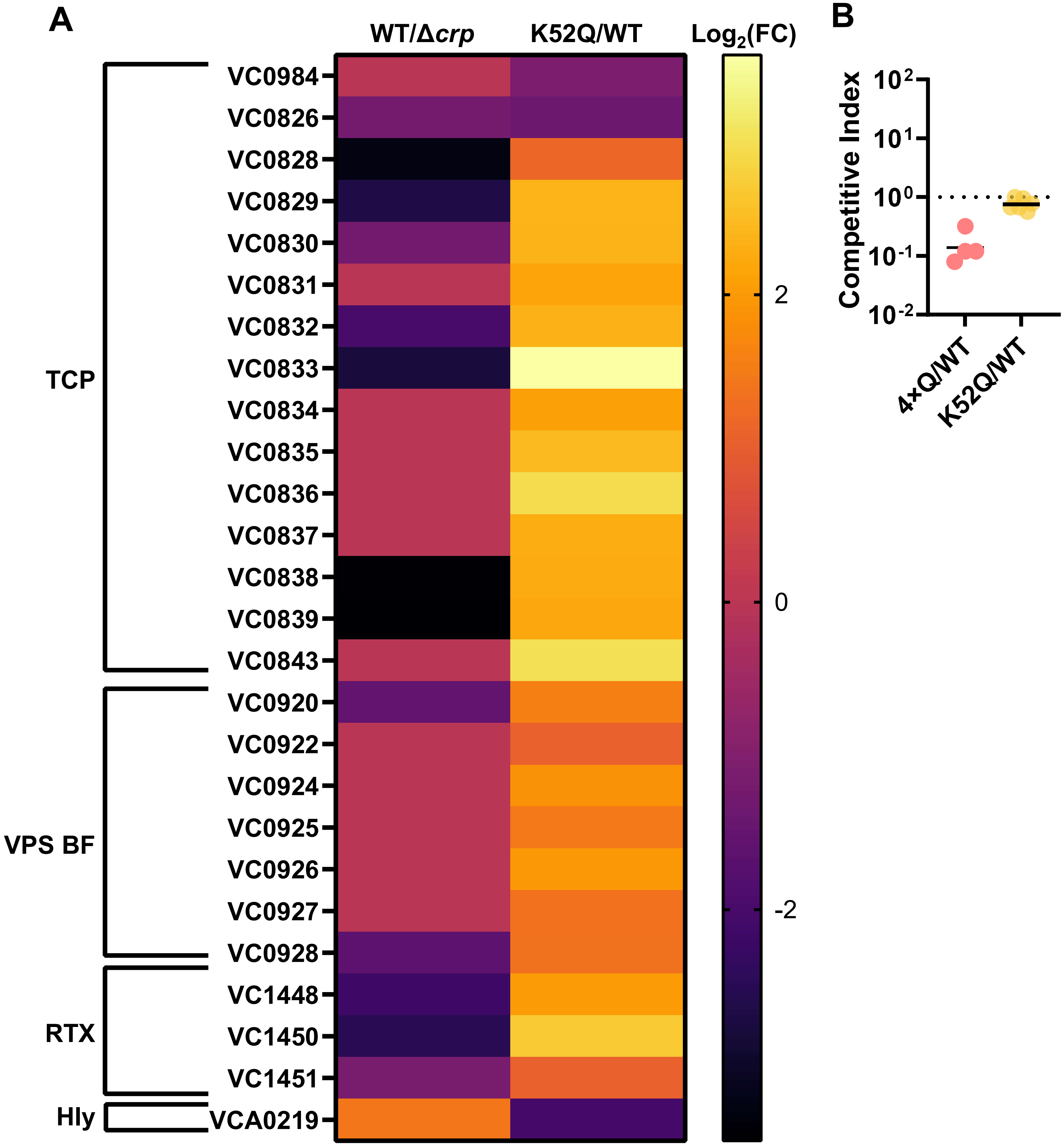


**Figure S6: The CRP K52Q mutant increases the transcript abundance of genes required for synthesis of the toxin co-regulated pilus, the VPS-dependent biofilm, and the RTX toxin.** Heat map illustrating the relative transcript abundance of the indicated genes in a K52Q mutant vs WT *V. cholerae.* EdgeR was used to identify significantly differently regulated genes Colors reflect the $\log_2$ (K52Q/WT transcript abundance). (B) Competitive index (CI) of the indicated strains in a neonatal mouse colonization model. CI was calculated as the output ratio of the V5-tagged variant allele to the WT control strain, normalized to the input ratio. The mean of at least 4 animals is shown. The two data sets are the product of independent experiments. Significance was calculated by applying an unpaired student's t test to log-transformed data. ** p<0.01, * p<0.05

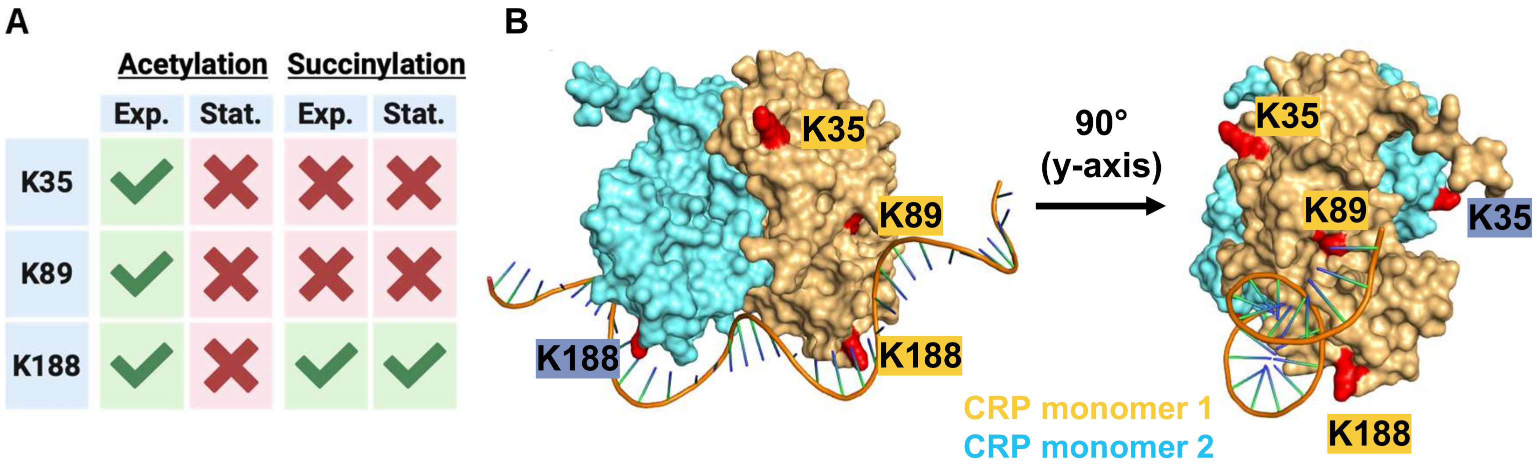


**Figure S7: In minimal medium supplemented with sucrose, *V. cholerae* CRP acetylation decreases in stationary phase.** (A) CRP-V5 affinity purification from exponential and stationary phase *V. cholerae* cultures followed by proteomic analysis for post-translational modifications. Acetylation and succinylation sites identified are shown. (B) Depiction of a CRP dimer bound to DNA with acetylated residues highlighted in red.

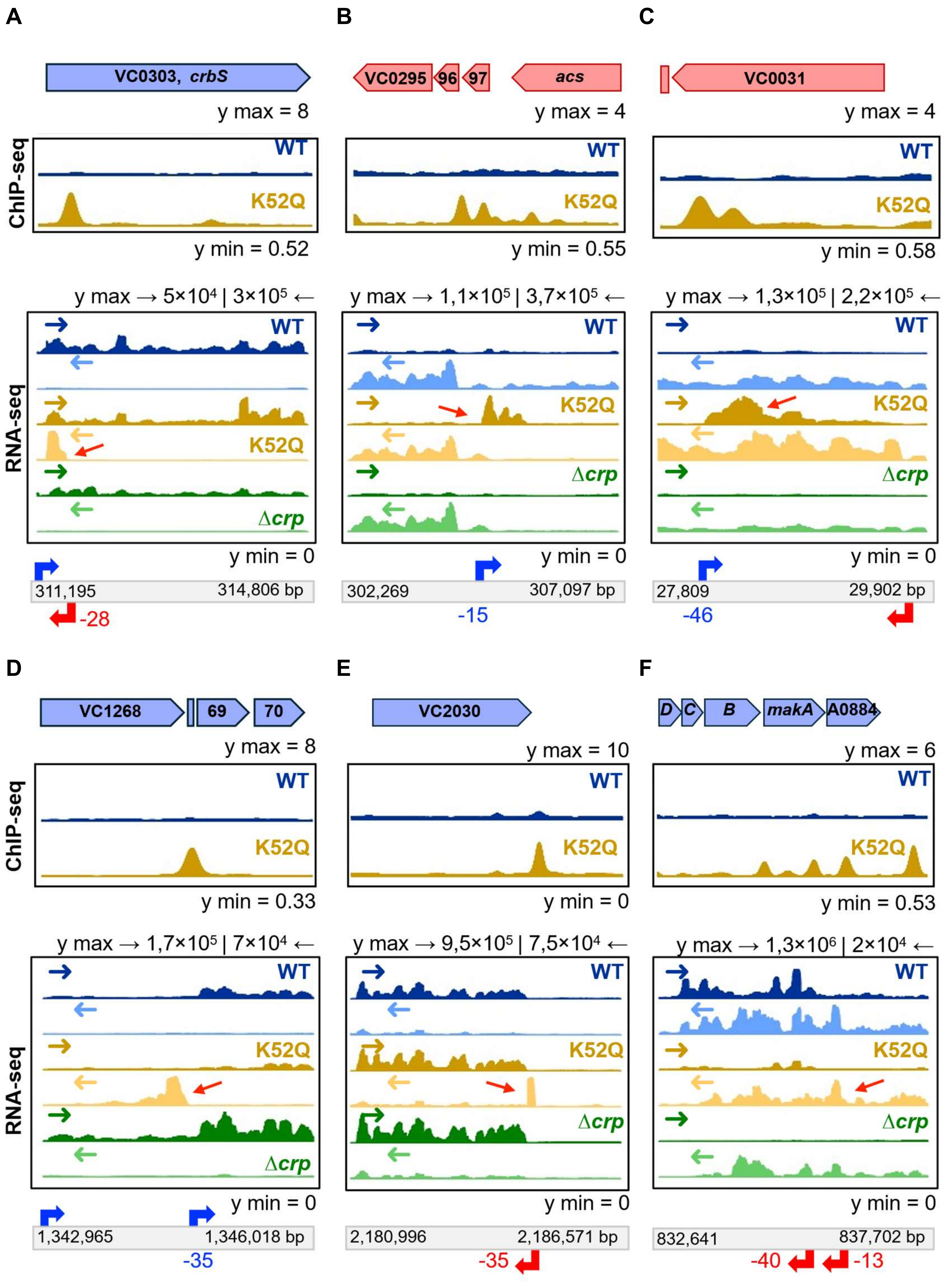

A
B
C
VC0303, crbS
VC0295
96
97
acs
VC0031
y max = 8
y max = 4
y max = 4
ChIP-seq
WT
K52Q
y min = 0.52
y min = 0.55
y min = 0.58
y max → 5×10⁴ | 3×10⁵ ←
y max → 1,1×10⁵ | 3,7×10⁵ ←
y max → 1,3×10⁵ | 2,2×10⁵ ←
RNA-seq
Δcrp
y min = 0
311,195
314,806 bp
302,269
307,097 bp
27,809
29,902 bp
-28
-15
-46
D
E
F
VC1268
69
70
VC2030
D
C
B
makA
A0884
y max = 8
y max = 10
y max = 6
y min = 0.33
y min = 0
y min = 0.53
y max → 1,7×10⁵ | 7×10⁴ ←
y max → 9,5×10⁵ | 7,5×10⁴ ←
y max → 1,3×10⁶ | 2×10⁴ ←
1,342,965
1,346,018 bp
2,180,996
2,186,571 bp
832,641
837,702 bp
-35
-35
-40
-13

**Figure S8: Six putative RNAs are uniquely activated by CRP K52Q.** ChIP-seq traces above (including an untagged CRP (Mock) and RNA-seq traces below of six putative RNAs whose transcript abundance is uniquely activated by CRP K52Q. Positions of the CDS for relevant genes are shown above. Arrows in each RNA-seq trace indicate sense (→) and antisense (←) directions. Bars below indicate genome coordinates shown. Arrows below indicate TSSs. Numbers below indicate the position of the CRP ChIP peak with respect to the TSS. (A) is also shown in Fig 5A.

**Genomic position. I: 311,200-311,700**

TCAAAAAACAACCTCGTTTTAATAGAAGCCTCTCAATCGAGAGGCTTTTTTCATTTCGC

AACAGCAAGG ATGTCTGATGCAAGGTTGGTTGGTAATTCCCGTTTCGCTCGCGTATCTCG

GCGTGCTGTTTTTGATTGCGTGGTATGGTGACAG GCAAACTCGTTGGCTCGCCAATTGGC

GGCCGTGGATCTATAGCTTGTCGATTGCGGTTTACTGT**ACCTCGTGGACGTTTTACGG**CA

CAGTCGGGCAGGCGAGTGTCAATCCTTGGTCGTTTCTGCCTATCTATATCGCCCCGATCT

TGATGTTTGTGT**TTGGTTGGCGAGTTTTAGCG**CGGCTGATTTTGATTGCTAAACGT GAGC

ACATTACTTCGCTCGCCGATTTTATTGCCGC CCGTTATGGTAAATCGCAGGGTTTGGCCG

**T**GGCGGTGACGGTGATCGCGGTCTTGGGTATTTTGCCGTATATCGCGCTACAACTGCGCG

GCATTACGATGGGATTGAACA

**Figure S9: Alignment of the circularized RNA sequence to the *V. cholerae* genome.** The genomic sequence above was extracted from *V. cholerae* chromosome I (bp 311,200 - 314,700 bp). cDNA generated from circularized RNA was amplified with the primers whose position and sequence are highlighted above in bold pink font. The arrows above the primer sequences indicate the direction of amplification. The sequence of the PCR product derived from CRP K52Q mutant RNA is shown in pink. The sequence of *crbS* not found in the PCR product is shown in red. A previously reported transcription start site located at 311,592 bp is indicated by the blue arrow, and the average maximum position of the CRP peak is shown in green.

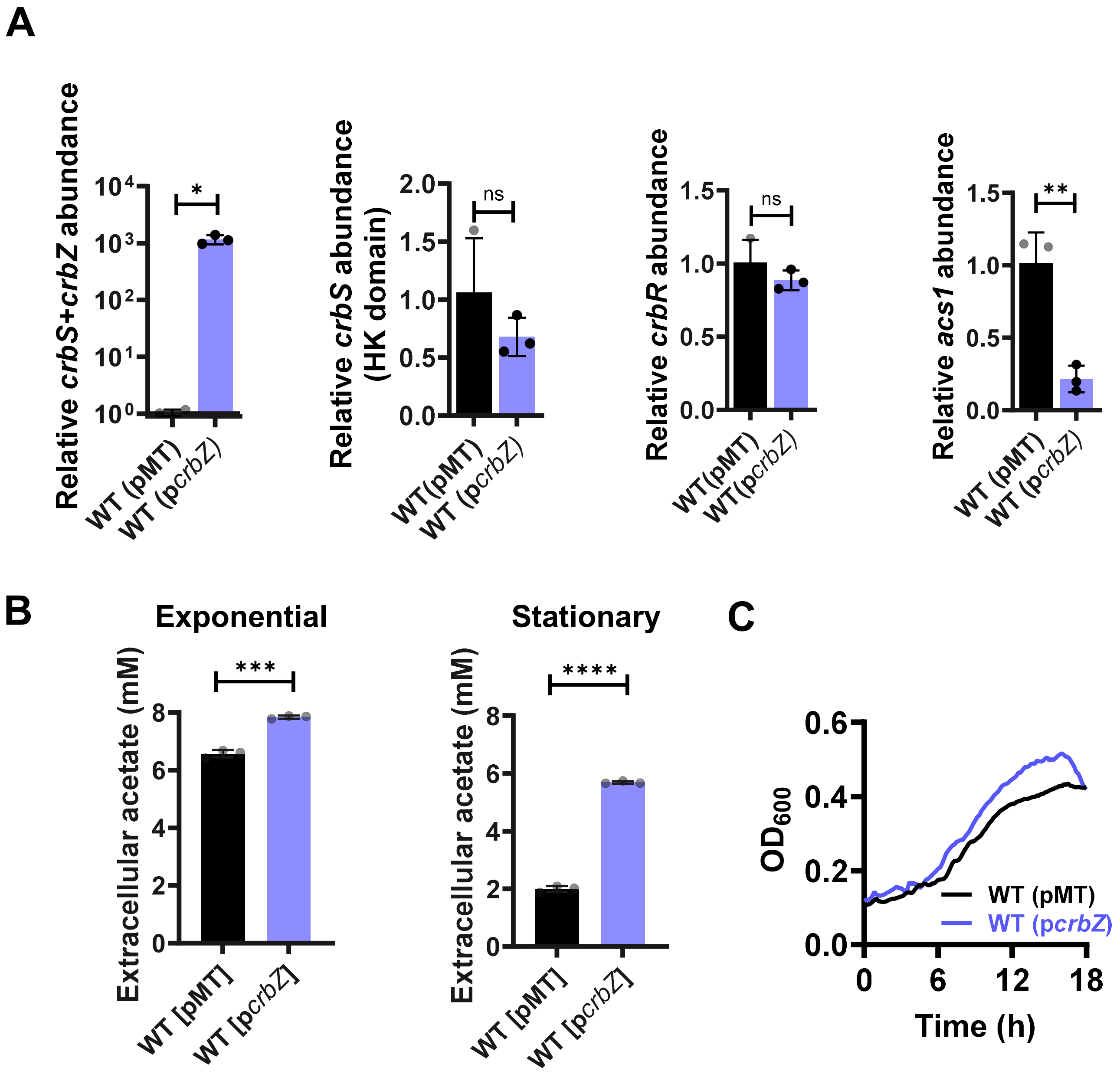


**Figure S10: The sRNA *crbZ* regulates the acetate switch and maltose metabolism.** (A) qRT-PCR analysis of the indicated genes in WT *V. cholerae* harboring either an empty vector (pMT) or a plasmid encoding *crbZ* from an inducible promoter. (B) Measurement of acetate in the supernatants of exponential and stationary phase LB cultures of the indicated strains. (C) Growth of the indicated strains over time in minimal medium supplemented with 0.4% maltose. The mean of biological triplicates is shown. Error bars represent the standard deviation. A student's t test was used to calculate significance. **** P<0.0001, *** p<0.001,** p<0.01, *p<0.05, ns not significant.

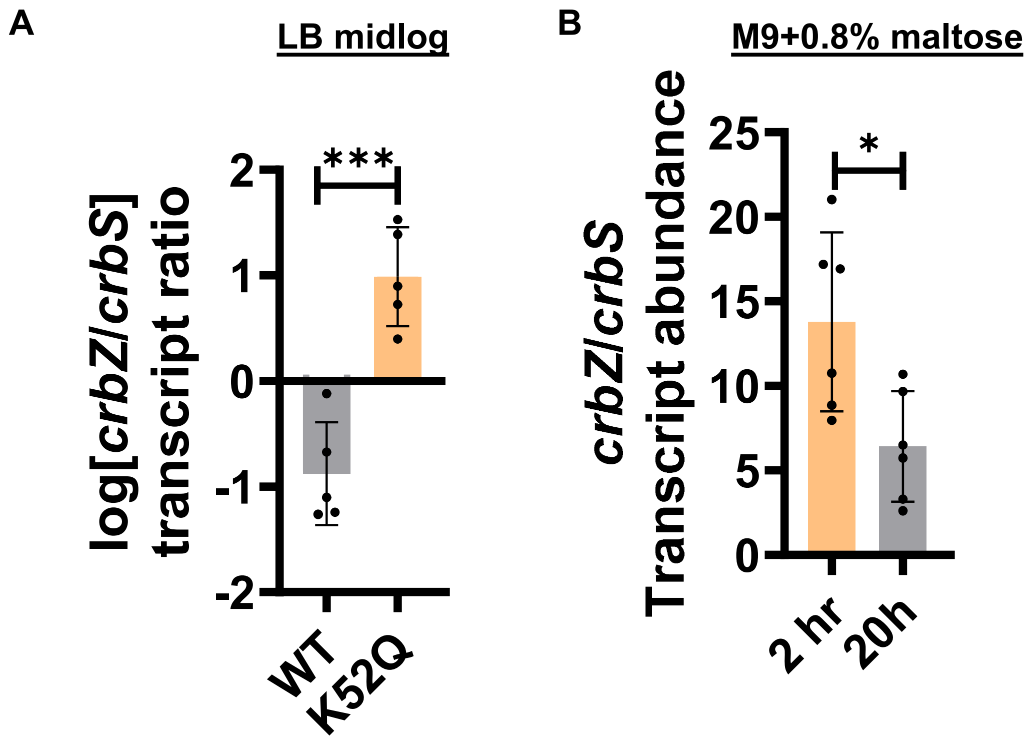


**Figure S11: Strand-specific RT-qPCR demonstrates activation of *crbZ* transcript abundance in maltose-containing medium.** Strand specific qRT-PCR of *crbZ* (anti-sense) and *crbS* (sense) for (A) WT *V. cholerae* or a CRP K52Q mutant cultured to mid-log phase in LB broth and (B) WT *V. cholerae* cultured for 2 or 20 h in M9 medium containing 0.8% maltose**.** Significance was calculated for (A) using a Welch's t test on log-transformed data and for (B) using an unpaired t test. The mean of at least 5 biological replicates is shown. *** P<0.001, * p<0.05.
