## Supplemental Methods for "Actuation of CRP activating region 3 by acetylation modulates *V. cholerae* sugar utilization and virulence"

### Supplementary Methods

**Cellular fractionation.** *V. cholerae* cellular fractionation protocol was performed as previously described (1). Briefly, mid-exponential-phase V5-tagged cells were lysed by sonication (Fisher Scientific, Sonic Dismembrator Model F60) in phosphate-buffered saline (PBS) supplemented with 1× protease inhibitor mixture (Boston Bioproducts, PI-210), 5 mM EDTA (Invitrogen 15575-038), and 1 mg/mL lysozyme (Alfa Aesar, J60701). Lysates were centrifuged to remove debris, and the resulting supernatant was filtered through a 0.22 µm syringe filter (VWR, 28145-501), followed by ultracentrifugation in a Beckman Coulter Optima L-90K Ultracentrifuge using a SW 41 Ti rotor at 30,000 rpm for 1h at 4 °C. The supernatant (cytoplasmic fraction) and the pellet (membrane-enriched fraction) were collected and stored at -20 °C until analysis. Fractionation efficiency was assessed by Western blot using an antibody against the RNA polymerase α-subunit antibody (BioLegend, 663104).

**Western blotting.** Cytoplasmic, membrane, and total cell lysates were prepared in 1× Laemmli buffer, separated by SDS-PAGE on 4-20% Mini-PROTEAN TGX gels (Bio-Rad, 4561096), and transferred to PVDF membrane (0.45 µm pore size) using the Trans-Blot Turbo Transfer System (Bio-Rad). V5-tagged proteins and RNAP α-subunit were detected using a monoclonal anti-V5 antibody (GenScript, A01724) at 1:2,000 dilution and an anti-RNAP α-subunit antibody (BioLegend, 663104) at 1:5,000 dilution. IRDye 800CW goat anti-mouse (LI-COR, 926-32210) and IRDye 680RD goat anti-rabbit secondary antibody (LI-COR, 926-86071) was used at a 1:2,000 dilution. Fluorescence signals were visualized using an Azure c600 imaging system (LI-COR) and quantified using ImageJ software.

**Immunoprecipitation and protein identification.** V5-tagged CRP was purified using the V5-Trap Magnetic Agarose Kit (Chromotek, V5TMAK-20) according to the manufacturer's instructions with minor adaptations. Briefly, 50 mL bacterial cultures were harvested and resuspended in dilution buffer (10 mM Tris pH 7.5, 150 mM NaCl, 0.5 mM EDTA) supplemented with protease inhibitor, DNaseI. After cell disruption by sonication using a Diagenode Bioruptor UCD-200

Sonicator (High Power setting; 30 s ON/ 30 s OFF, four cycles of 30 s each), clarified lysates (400 µL) were incubated with 50 µL of equilibrated V5-trap beads overnight at 4 ˚C with gentle rotation. Beads were washed five times (dilution buffer supplemented with 0.05% Triton X-100), and eluted in 80 µL of Laemmli buffer at 95 ˚C for 10 min. Eluted proteins were resolved by SDS-PAGE (4-20% TGX gels, Bio-Rad) and stained with Imperial Protein Stain (Thermo Scientific). Bands near the 25 kDa were excised and submitted to the Taplin Mass Spectrometry Facility (Harvard Medical School) for LC-MS/MS analysis to identify CRP and assess post-translational modifications, including lysine acetylation and succinylation. Predicted structural models of *V. cholerae* CRP were generated using AlphaFold3 with the reference sequence from UniProt (Q9KNW6) to highlight the modified residues.

**Growth curves.** For experiments started from an LB broth pre-culture, cells were grown to mid-exponential phase in LB broth and then diluted 1:100 into the indicated medium. Cells were washed prior to dilution into minimal medium. For pre-cultures in M9 minimal medium supplemented with 0.4% maltose, cells were grown to an $0D_{600}$ of 0.05 and inoculated at a 1:10 dilution into fresh medium with the indicated concentration of maltose. Bacterial growth was monitored in Nunc Edge 96-well flat-bottom microplates (Thermo Fisher). Each well contained 200 µL of culture. The peripheral edge wells were filled with sterile phosphate-buffered saline (PBS) to minimize evaporation. Plates were incubated at room temperature, and the $OD_{600}$ was recorded every 30 min for 18 h using SpectraMax ABS or SpectraMax iD5 microplate readers (Molecular Devices).

**Virulence assays.** All animal experiments were conducted in accordance with protocols approved (IS00000116-9) by the Harvard Medical School Institutional Animal Care and Use Committee (IACUC), adhering to NIH guidelines. *V. cholerae* infections of suckling mice were performed as previously described (2). In brief, five-day-old infant mice were separated from their mothers and inoculated with ~5x$10^6$ colony-forming units (CFUs) of total bacteria suspended in LB media. In these competition experiments, a 1:1 mixture of two strains was administered via feeding tube into the stomach using a 50 µL inoculum. Mice were sacrificed after 22 hours, and the small intestines were homogenized immediately in 5ml LB media using a Tissue-Tearor™ (BioSpec 985370). Enumeration of each strain in the starting inoculum and those recovered from the small intestine at the endpoint was performed by selective plating of serial dilutions on LB 1.6% agar plates supplemented with Streptomycin (200 µg/ml) and X-Gal (20 mg/ml).

**Strand-specific reverse transcription (RT).** For strand-specific transcript detection, reverse transcription was performed using gene specific primers complementary to either the sense or antisense strand, as listed in Supplementary Table S5. Equal amounts of total RNA (500 ng) were used in each reaction, and cDNA synthesis was carried out using SuperScript III First-Strand Synthesis System (Invitrogen). Quantitative PCR was performed as previously described. Transcript abundance was calculated using ∆Ct values derived directly from Cq measurements. Comparisons were made only between samples generated under identical conditions. At least five independent biological replicates were used.


1. J. A. Gibson, M. J. Gebhardt, R. Santos, S. L. Dove, P. I. Watnick, Sequestration of a dual function DNA-binding protein by Vibrio cholerae CRP. *Proceedings of the National Academy of Sciences of the United States of America* **119**, e2210115119 (2022).

2. M. J. Angelichio, J. Spector, M. K. Waldor, A. Camilli, Vibrio cholerae intestinal population dynamics in the suckling mouse model of infection. *Infection and immunity* **67**, 3733–3739 (1999).
